# Lineage specifying transcription factors determine cell function by direct control of metabolism

**DOI:** 10.64898/2026.07.02.735862

**Authors:** Alex J. Bott, Joshua C. Youngs, Corey N. Cunningham, Jonathan G. Van Vranken, Peng Wei, Ahmad A. Cluntun, Steven P. Gygi, Donald E. Ayer, Jared Rutter

**Affiliations:** Department of Biochemistry, University of Utah, Salt Lake City, United States; Department of Cell Biology, Harvard Medical School, Boston, United States; Department of Oncological Sciences, Huntsman Cancer Institute, University of Utah, Salt Lake City, United States; Howard Hughes Medical Institute, University of Utah, Salt Lake City, United States

**Author notes:** Department of Biochemistry and Molecular Biology, Rutgers University, Piscataway, United States (Ahmad A. Cluntun).

## Abstract

Execution of a complex cellular function requires the coherent expression of a set of genes that encode the requisite biochemical program. It is believed that this is typically accomplished by employing common transcription factors to activate those genes, thus ensuring coherence. However, many of these programmatic gene sets are expressed at differing levels and in distinct configurations in different cell types. This creates a unique challenge for the logic of gene regulation—precisely tuning the expression of programs across cell types while maintaining coherence. Using the glycolysis pathway as a model of co-expressed genes with highly divergent cell-specific regulation, we developed a CRISPR screening approach to identify trans-acting factors that control RNA abundance. Using this platform, we discovered that lineage-specifying transcription factors directly activate glycolytic gene expression and alter metabolism. We showed that impaired metabolism disrupts lineage-specific activities via loss of specialized biochemical pathways and functions. We hypothesize that lineage-specifying transcription factors directly regulate metabolic gene expression to ensure that cellular metabolism is tuned to the specific demands imparted by specialized cell function.

## Introduction

Cellular specialization requires different cell types to execute distinct biochemical programs, yet many of the pathways underlying these programs, including glycolysis, are expressed in virtually every cell in the body. Glycolysis is an ancient pathway, conserved across all kingdoms of life, by which carbohydrates are assimilated for energy extraction and generation of biosynthetic building blocks (*1, 2*). Because each enzyme in the cascade accepts the product of the upstream step as its substrate, a gene encoding every step in the pathway must be expressed to complete the metabolic route, creating a strong selective pressure for coherent regulation. Consistent with this, glycolytic genes are induced coordinately in response to proliferative signals via MYC or hypoxia via HIF1α. Yet despite this coherent inducibility, the expression level of each glycolytic step, and which isoenzyme executes it, is cell-type-specific. How different cell types can each maintain a coherent glycolytic program while tuning its configuration to their own unique metabolic demands is a fundamental and unresolved question in gene regulation.

One approach to this question is to identify the transcription factors that drive glycolytic gene expression in each cell type. While it is relatively straightforward to identify the targets of a given transcription factor, determining which transcription factors regulate a specific gene of interest is substantially harder. The tools to systematically do so, at endogenous loci, and in a manner broadly accessible, remain limited. Therefore, we developed Targeted Readout to Understand Transcription via Fluorescent In Situ Hybridization (TRoUT-FISH), a CRISPR-based screening platform that identifies trans-acting regulators of any target RNA. Deploying TRoUT-FISH across glycolytic genes in multiple cell types, we discovered that lineage-specifying transcription factors directly activate glycolytic gene expression and alter metabolism. We show that this regulation is direct, that it is a general property of lineage-specifying transcription factors across multiple lineages, and that disrupting the resulting metabolic program impairs the specialized biochemical functions that define each lineage.

## Results

### TRoUT-FISH enables discovery of gene regulators

Artificial reporter constructs have been employed to discover regulators of gene expression, however, many aspects of gene regulation are only operative in the native genomic context (*3, 4*). Inspired by efforts to causally link non-coding regulatory elements to gene expression (*5*–*7*), we developed TRoUT-FISH, a screening platform to identify genes that impact the abundance of a specific target RNA (Fig. 1A). TRoUT-FISH harnesses nucleic acid chemistry to design highly sensitive and specific RNA FISH probes, thereby avoiding the use of either exogenous reporters or endogenous genome modification—both of which are time consuming, subject to artifacts, and difficult in certain cellular scenarios (*8*). Cells are perturbed with an sgRNA library and sorted by flow cytometry according to target RNA abundance as assessed by fluorescence. Those sgRNAs that are enriched in the sorted populations then identify putative activators and repressors of mRNA abundance.

**Figure 1:**
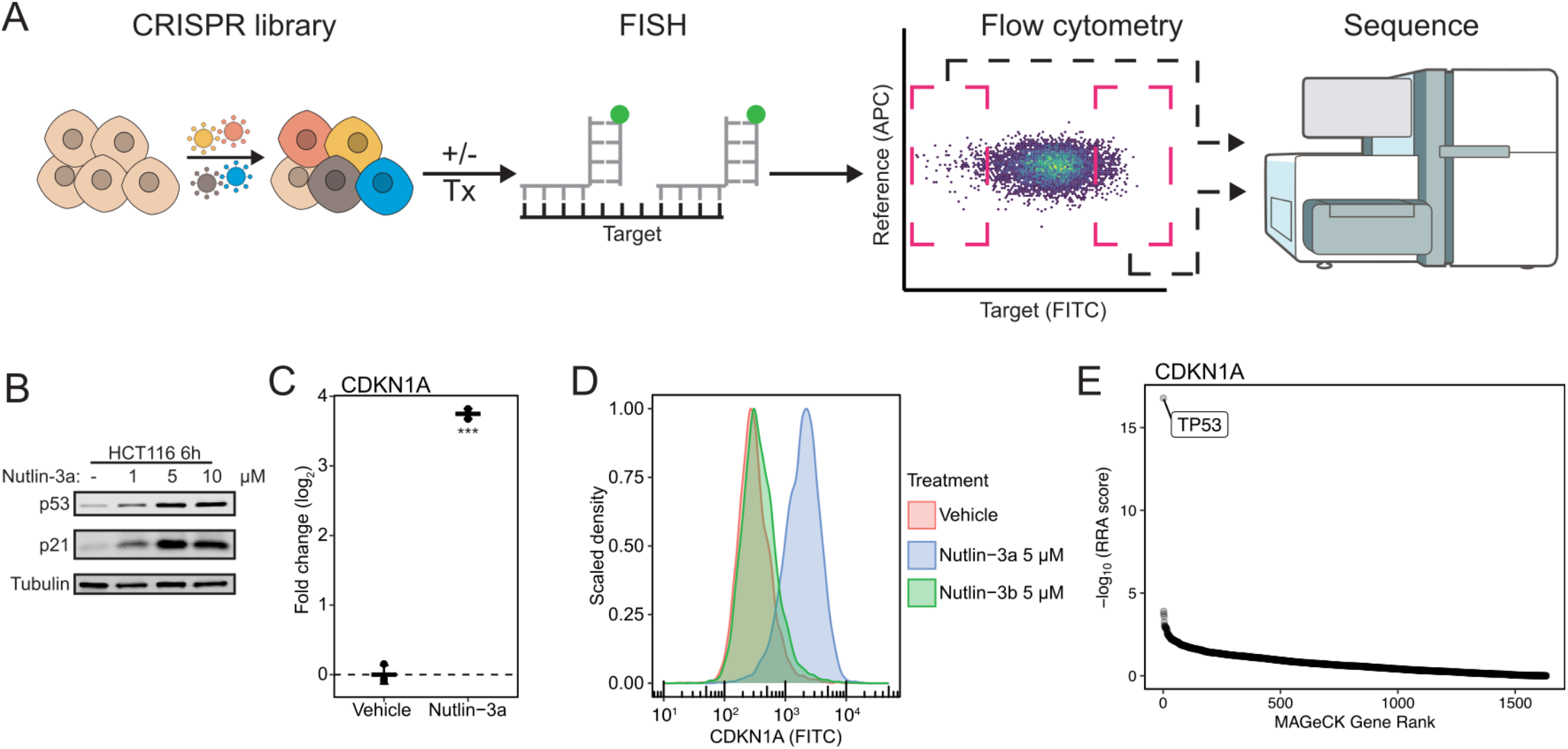
TRoUT-FISH enables discovery of gene regulation. **(A)** Schematic of TRoUT-FISH. Cells are transduced with a pooled CRISPR library and selected (left), optionally exposed to a stimulus, and fixed. Cells undergo RNA FISH (middle-left) for targets of interest and sorted by flow cytometry (middle-right) based on expression of RNA targets, and sgRNA abundance is determined by sequencing (right). **(B)** Immunoblots of HCT116 cells treated with Nutlin-3a for 6 hours at the indicated doses. **(C)** Dot plot showing changes in CDKN1A expression in HCT116 cells treated with 5 µM Nutlin-3a for 6 hours, as determined by qPCR. Experiments performed in triplicate and means ± SD is shown. A two-tailed, two-sample t-test was performed (\*\*\**P* < 0.001). **(D)** RNA FISH and flow cytometry of CDKN1A of HCT116 cells were treated with 5 µM Nutlin-3a or Nutlin-3b for 6 hours. Experiments performed in triplicate. **(E)** TRoUT-FISH results for CDKN1A abundance in HCT116 cells treated with Nutlin-3a. RRA scores derived from MAGeCK comparing the high and low populations of CDKN1A abundance of Nutlin-3a treated cells.

To test the effectiveness of TRoUT-FISH, we turned to a well-defined paradigm of drug-induced transcriptional regulation with a known mediator. The small molecule Nutlin-3a inhibits the interaction between MDM2 and p53, thus preventing ubiquitination and degradation of p53 (*9, 10*), enabling it to promote expression of target genes such as *CDKN1A* (Fig. S1A). We treated *TP53* (p53) wild-type cells with Nutlin-3a and observed a robust increase in p53 protein and induction of both *CDKN1A* transcript and the p21 protein it encodes (Fig. 1B,C, S1B). Using FISH probes tiled across *CDKN1A* exons (Fig. S1C) (*11*–*13*), we observed an increase in *CDKN1A* FISH signal intensity in cells treated with Nutlin-3a relative to vehicle or the inactivate enantiomer (Nutlin-3b), suggesting reliable quantification of transcript abundance via FISH and flow cytometry (Fig. 1D). We then performed TRoUT-FISH with an sgRNA library targeting transcription factors (*14*) to identify genes affecting *CDKN1A* mRNA abundance. Cells were treated with vehicle or Nutlin-3a, subjected to *CDKN1A* RNA FISH, and *CDKN1A*^low^ and *CDKN1A*^high^ populations were sorted and sgRNAs sequenced. The one gene that was clearly required for Nutlin-3a-induced *CDKN1A* expression was *TP53* (Fig. 1E, S1D). As validation, we generated stable *TP53* knockout lines (Fig. S1E) and observed no induction of *CDKN1A* FISH signal upon treatment with Nutlin-3a (Fig. S1F, G), consistent with published data (*15*).

These data suggest that TRoUT-FISH identifies transcription factors that are required for stimulus-responsive gene expression. To test the ability of TRoUT-FISH to identify the mechanisms underlying basal transcriptional regulation, we designed probes against *TXNIP* (Fig. S2A), which is regulated by media glucose concentration (*16*). We cultured cells in 0 or 11.1 mM glucose for 16h, added glucose back to one culture for 6h, then measured *TXNIP* abundance by FISH. Glucose withdrawal decreased *TXNIP* levels to near background with glucose addback rescuing expression, though not fully to replete levels (Fig. S2B). Having confidence in the fidelity of the *TXNIP* FISH signal, we performed a TRoUT-FISH screen with an epigenetic-focused CRISPR library (*17*). We observed *MLX* and *MLXIP* among the top three putative *TXNIP* activators (Fig. S2C-E). *MLX* and *MLXIP* are basic helix-loop-helix transcription factors that heterodimerize and are the best-characterized activators of *TXNIP* expression (*16, 18, 19*). Together, these data suggest that TRoUT-FISH can discover genes required for either basal or stimulated transcriptional regulation.

### Transcriptional control of glycolytic genes

Having established the effectiveness of TRoUT-FISH to discover gene regulatory factors, we proceeded to interrogate the transcriptional control of glycolysis (Fig. 2A). To identify a cellular model in which most glycolytic genes were expressed at reliably measurable levels, we queried DepMap gene expression data for >1000 cell lines (*20*). We extracted expression levels for glycolytic genes and scored based on cumulative expression rank. The top one hundred scoring cell lines were enriched for cells from either skin or central nervous system lineages (Fig. S3A,B), which might reflect metabolic features of these cancers (*21*–*23*).

**Figure 2:**
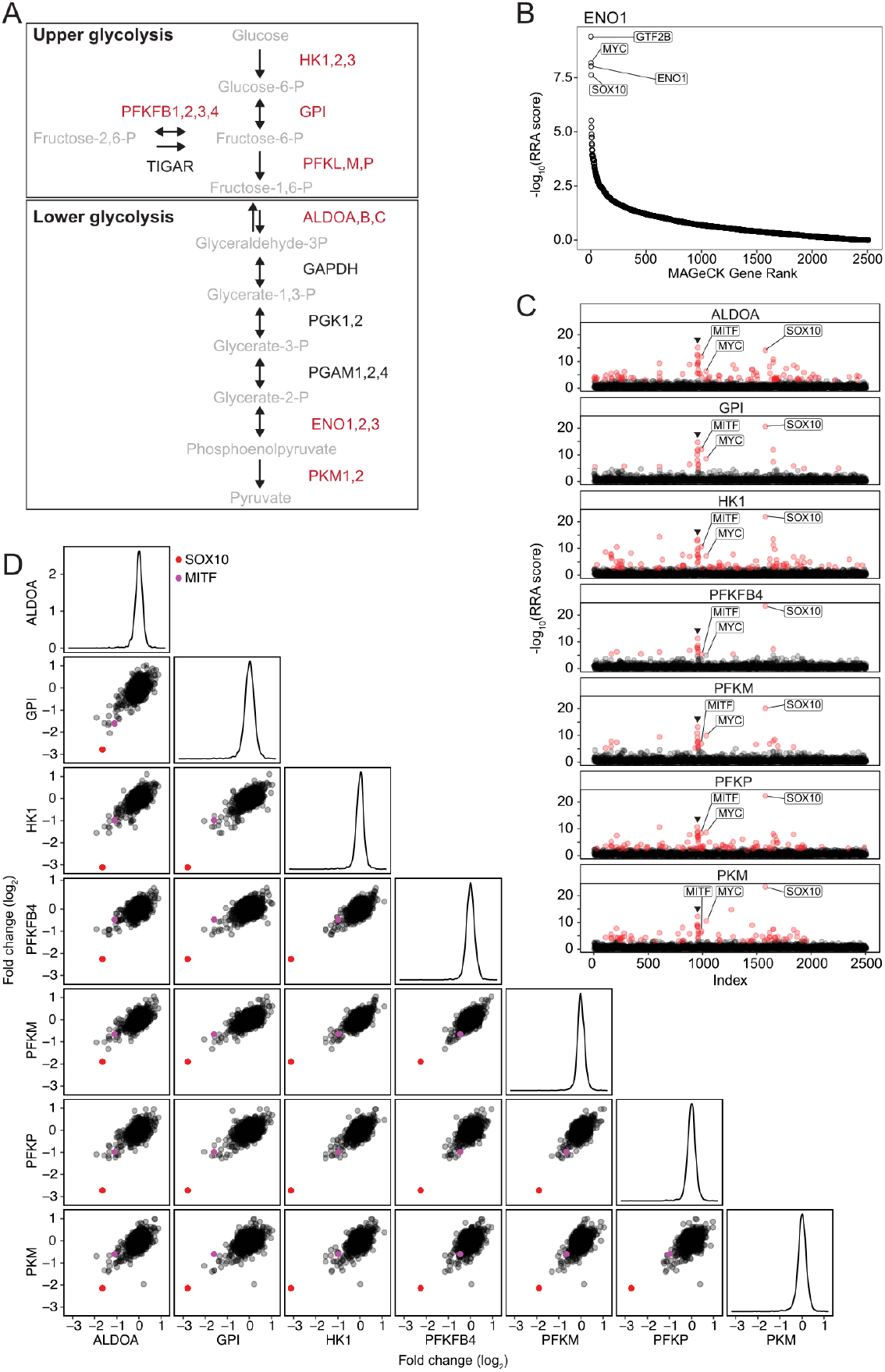
TRoUT-FISH identifies the lineage-specifying transcription factors *SOX10* and *MITF* as regulators of glycolytic gene expression in melanoma cells. **(A)** Schematic of the glycolytic pathway. Metabolites in gray, genes in black, and genes measured by TRoUT-FISH in MeWo cells in red. **(B)** TRoUT-FISH identifies regulators of ENO1 in MeWo cells. RRA scores derived from MAGeCK comparing the high and low populations of ENO1 abundance. **(C)** TRoUT-FISH identifies regulators of glycolytic target genes in MeWo cells. Scatterplots show genes indexed alphabetically along the x-axis and the RRA scores from MAGeCK comparing the high and low populations of each target gene along the y-axis. Genes with FDR ≤ 0.05 are colored red. Mediator genes indicated by black arrowhead. **(D)** TRoUT-FISH results across glycolytic genes. Pairwise scatterplots display the gene level log_2_ fold change comparing the high and low populations for each target gene. On the diagonal are density plots displaying the distribution of gene level effects. Red is SOX10 and magenta is MITF in each scatterplot.

We performed TRoUT-FISH screens for several glycolytic genes in MeWo cells, which are *BRAF* WT melanoma cells that express most glycolytic genes to a degree detectable by FISH (Fig. S3C orange dot; Fig. S4A) and lack significant glycolytic gene copy number changes (Fig. S3D). We first performed TRoUT-FISH using an epigenetic focused CRISPR library (*17*) for *ENO1*, the product of which catalyzes the conversion of glycerate-2-phosphate to phosphoenolpyruvate (Fig. 2B). Satisfyingly, TRoUT-FISH identified *ENO1* sgRNAs (included in this library as the short form was shown to transcriptionally repress *MYC* (*24*)) as strongly depleting *ENO1* RNA, serving as an internal positive control (Fig. 2B, S4C). Our TRoUT-FISH screen also identified *MYC* as a positive regulator of *ENO1* as has been previously reported (Fig. 2B, S4B). Among other high scoring candidate transcriptional regulators were the general transcription factor *GTF2B* and *SOX10*, (Fig. 2B, S4D). We proceeded to perform TRoUT-FISH for seven additional glycolytic genes in MeWo cells using the same epigenetic library. Target genes returned a variety of putative regulators, with the genes encoding subunits of the Mediator complex being found universally (Fig. 2C – arrowhead). However, the top scoring positive regulator was *SOX10* for all genes, with *MITF* also being a highly significant hit (Fig. 2D – red and magenta dots, respectively). These data, along with our observation that the skin lineage was amongst the most enriched for high glycolytic gene expression, suggest that the lineage-specifying transcription factors (LSTFs) *SOX10* and *MITF* drive a glycolytic metabolic gene expression program alongside their canonical differentiation functions.

### *SOX10* and *MITF* directly regulate glycolytic genes

To further test the results of the TRoUT-FISH screens, we generated MeWo cells expressing nuclease-dead Cas9 fused to KRAB and MeCP2 (hereafter called MeWo_KRAB), allowing site specific recruitment of repressive machinery to transcription start sites (*25*–*27*). MeWo_KRAB cells infected with 2sgRNAs each against *MITF* or *SOX10* showed decreased transcript for their respective targets. In addition, silencing of *SOX10* reduced *MITF* expression which is known to be positively regulated by *SOX10* (Fig. S5A). To understand the effects of *SOX10* and *MITF* suppression transcriptome-wide, we performed RNA-sequencing and found that replicates, sgRNAs, and genotypes clustered tightly (Fig. S5B). We observed the hierarchical control of *MITF* by *SOX10* in our sequencing data (Fig. 3A). In addition, silencing either of these LSTFs decreased the expression of the canonical structural genes of melanogenesis, as expected (Fig. 3B,C). It also decreased the expression of glycolytic genes, consistent with our TRoUT-FISH results (*28*). Silencing either LSTF altered metabolic gene expression more broadly, suggesting that these factors, either directly or through downstream effectors, drive a particular metabolic program (Fig. S5C,D). To test whether this regulation is direct, we performed ChIP-sequencing for MITF. As expected, we observed MITF peaks at canonical melanogenesis genes (Fig. 3D, S5E). We also found the MITF bound directly to glycolytic genes (Fig. 3E, S5F), suggesting that the relationship between MITF and glycolytic genes is direct.

**Figure 3:**
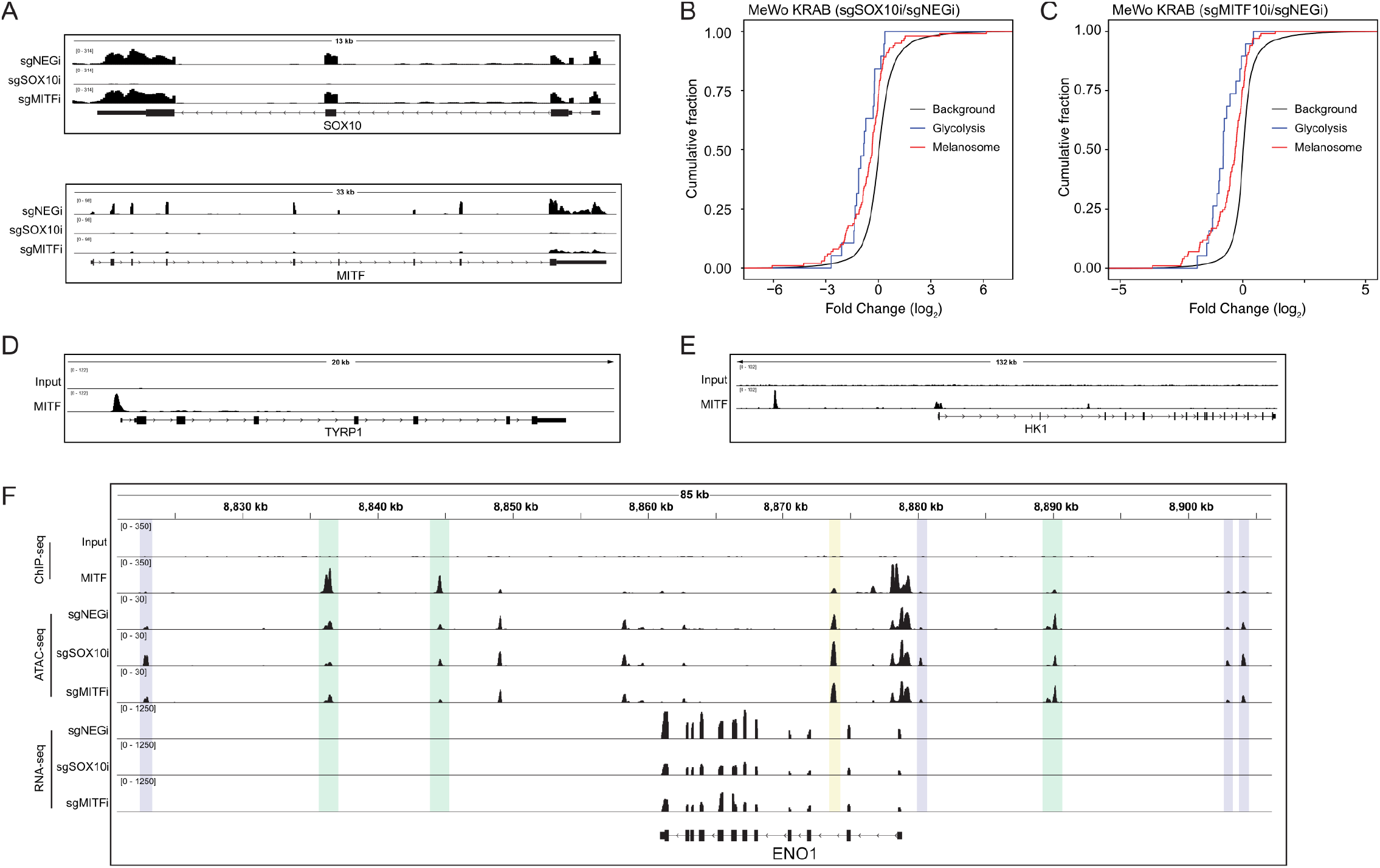
*SOX10* and *MITF* control glycolytic gene expression. **(A)** Genome browser tracks visualizing of RNA-sequencing in MeWo KRAB cells. **(B)** Cumulative distribution plots displaying log2 fold change comparing SOX10 silenced cells to control. Genes are annotated as related to glycolysis, melanosome, or background set. **(C)** Cumulative distribution plots displaying log2 fold change comparing MITF silenced cells to control. Genes are annotated as related to glycolysis, melanosome, or background set. A two-tailed, two-sample Kolmogorov–Smirnov test was performed between glycolysis and background annotation (*P* = 1.122 × 10^−4^). **(D)** Genome browser tracks visualizing of ChIP-sequencing peaks of MITF binding in MeWo cells near canonical melanosome related genes. A two-tailed, two-sample Kolmogorov– Smirnov test was performed between glycolysis and background annotation (*P* = 2.747 × 10^−7^). **(E)** As in (D), but near glycolytic genes. **(F)** Genome browser tracks visualizing MITF ChIP-sequencing, ATAC-sequencing, and RNA-sequencing in MeWo_KRAB cells. The purple regions are increased accessibility with a RUNX and/or HAND motif, the yellow region is increased accessibility with AP1/CEBPE/Maz motif, and the green regions are decreased accessibility for either or both genotypes.

Recent studies have shown that many LSTFs alter chromatin accessibility as part of the mechanism whereby they induce the expression of target genes (*29*–*31*). To assess whether this is true of SOX10 and MITF on glycolytic genes, we silenced their expression and performed ATAC-sequencing. As before, replicates, sgRNAs, and genotypes clustered tightly (Fig. S6A). We performed *de novo* motif discovery of differentially accessible regions (*32*). Silencing *SOX10* or *MITF* led to loss of both motifs (Fig. S6B,C) and substantial changes to chromatin accessibility, including regions proximal to metabolic genes (Fig. S7A-H). Suppressing either LSTF decreased chromatin accessibility at *MITF* binding sites proximal to canonical melanogenesis genes (Fig. S6D, green). It also decreased chromatin accessibility at glycolytic genes, including *ENO1* (Fig. 3F, green). Taken together, these data show that SOX10 and MITF drive metabolic gene expression and that MITF directly binds the genome proximal to glycolytic genes.

### Inhibition of central carbon metabolism disrupts melanosomes

Our data showed that suppressing *SOX10* or *MITF* in melanoma cells reduced expression of glycolytic genes. To test whether this has a measurable impact on metabolism, we cultured MeWo *SOX10* CRISPRi cells with ^13^C-glucose and measured labeled metabolites by LC-MS. Sampling the media revealed less M+3 labeled lactate was secreted (Fig. 4A). Measuring intracellular metabolites showed that *SOX10* silenced cells had similar relative glucose abundance compared to controls but exhibited less labeling of glycolytic and TCA cycle metabolites (Fig. 4B). This suggests that, indeed, *SOX10* and *MITF* are required to maintain normal glycolysis.

**Figure 4:**
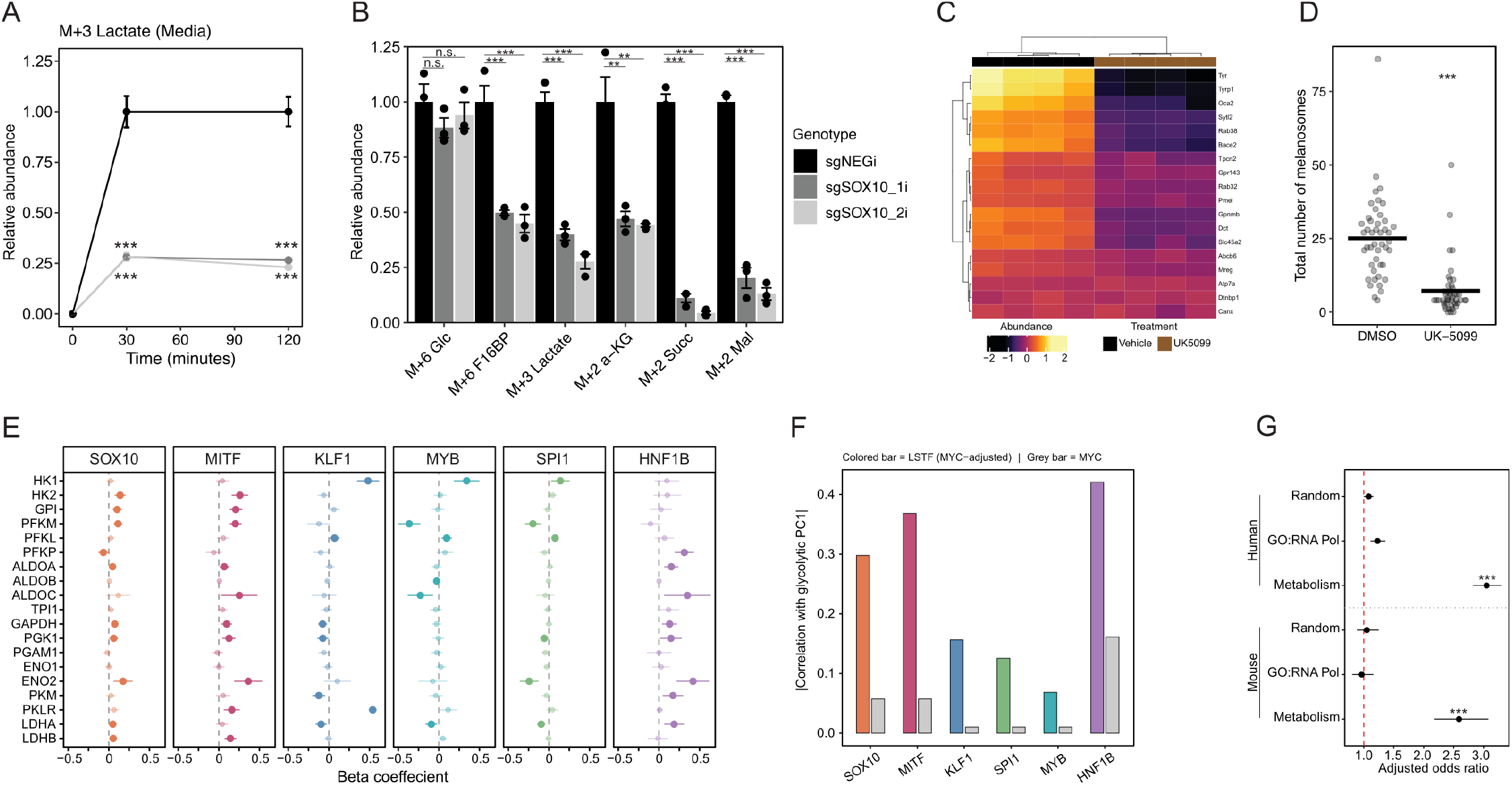
Inhibition of central carbon metabolism disrupts melanogenic processes. **(A)** Quantification of ^13^C-glucose tracing to M+3 lactate secreted from MeWo_KRAB cells transduced with guides against SOX10. A one-way ANOVA and Dunnett test were performed (\**P* < 0.05, \*\**P* < 0.01, \*\*\**P* < 0.001). **(B)** Quantification of ^13^C-glucose tracing (30 minutes) to indicated metabolites from MeWo_KRAB cells transduced with guides against SOX10. A one-way ANOVA and Dunnett test were performed (\**P* < 0.05, \*\**P* < 0.01, \*\*\**P* < 0.001). **(C)** Heatmap depicting change in melanosome proteins from B16F10 cells treated with vehicle or 10 µM UK-5099 for 48 hours. **(D)** Quantification of the number of total melanosomes measured by TEM. A two-tailed, two-sample t-test was performed (\**P* < 0.05, \*\**P* < 0.01, \*\*\**P* < 0.001). **(E)** Beta coefficient of intra-lineage ordinary least-squares regression of each glycolytic gene against LSTF expression controlling for *MYC* expression. Opaque points with larger symbols indicate genes passing FDR < 0.10 (Benjamini–Hochberg correction within each LSTF); translucent points are non-significant. Error bars represent 95% confidence intervals. **(F)** Absolute partial correlation (|partial *r*|) between each LSTF and the intra-lineage glycolytic PC1, after removing the linear contribution of *MYC* from both variables (colored bars). Absolute Pearson correlation between *MYC* and glycolytic PC1 (gray bars). Adjusted odds ratio of lineage-specific transcription factor binding to gene set transcriptional start site (+/-5kb) across human (n = 32730) or mouse (n = 22745) ChIP-sequencing datasets (\*\*\**P* < 0.001). Error bars represent 95% confidence intervals, dashed red line indicates OR = 1.

The mitochondrial pyruvate carrier (MPC), an obligate heterodimer of MPC1 and MPC2 that imports the product of glycolysis into the mitochondrial matrix where it can be oxidized, has been shown to impact cellular fate (*33*– *38*). We treated B16F10 mouse melanoma cells with the well-characterized inhibitor of the MPC, UK-5099, and observed a decrease in the cellular melanin content (Fig. S8C) without impacting cell proliferation (Fig. S8A). UK-5099 treatment also decreased the PMEL protein, a resident structural protein of the melanosome and target of *SOX10* and *MITF* (Fig. S8B). To gain a systematic understanding of cellular changes in response to UK-5099, we used tandem mass tag (TMT)-based quantitative proteomics. GO term enrichment for biological processes of downregulated proteins revealed melanin biosynthetic processes as the most enriched, with various melanin and pigment GO terms in the top ten (Fig. S8D). More focused analysis showed that cells treated with UK-5099 exhibited lower abundance of melanosomal proteins (Fig. 4D, S8E,F). This decrease in melanosomal protein abundance led us to hypothesize that disrupting central carbon metabolism may impact melanosomes themselves. Indeed, transmission electron microscopy (TEM) revealed marked defects in melanosomes (Fig. S8G,H) with decreases in the total number of melanosomes (Fig. 4D) and those in the late stage of assembly (Fig. S8I).

This connection between glycolytic gene expression and melanogenesis was also observed in RNA-sequencing data from human pluripotent stem cells progressively differentiated to neural crest, melanoblasts, or melanocytes (*39*), where glycolytic gene expression is higher in the fully, terminally differentiated melanocytes compared to neural crest cells (Fig. S9A). During our routine culture of B16F10 cells, we noticed that cells became less pigmented at higher passage number (Fig. S9B). RNA-sequencing of early and late B16F10 cells revealed coordinated decreases in both melanosome and glycolytic gene expression in late passage cells (Fig. S9C), suggesting that this correlation might also hold in this scenario. Together, these data show that glycolytic gene expression increases progressively during differentiation to melanocytes but suppressing *SOX10* decreases glycolytic flux and impairs specialized processes such as pigment synthesis. This model suggests that *SOX10* and *MITF* control metabolism, in part, to support the unique biochemistry that occurs in melanocytes.

### Lineage-specific transcription factors generally bind glycolytic genes

Given these observations, we sought to test whether LSTF(s) might directly regulate glycolytic genes in another lineage. K562 cells are erythroleukemic cells derived from a patient with chronic myelogenous leukemia (*40*). In our analysis, K562 cells did not exhibit copy number amplification of glycolytic genes and exhibited measurable expression of most glycolytic genes (Fig. S3C,D blue dot). We performed TRoUT-FISH for seven genes in K562 cells utilizing the same epigenetic CRISPR library (Fig. S10). We again found lineage-specifying transcription factors as transcriptional activators of glycolysis. *KLF1* and *NFE2* were identified as positive regulators of the expression of all glycolytic genes, and *MYB* and *GATA1* were identified as positive regulators of a subset of them (Fig. S10A).

As with MeWo cells, TRoUT-FISH identified *MYC* as a positive regulator of glycolytic gene expression in K562 cells. For several target genes, *BRD4*, a known regulator of *MYC*, was also identified as a positive regulator. We treated K562 cells with JQ1, which downregulates *MYC* transcription via inhibition of BRD4 (*41*), and observed decreased MYC transcript and protein (Fig. S11A,B). We also observed decreased expression of glycolytic genes to varying degrees (Fig. S11C). To directly test whether *MYC* or LSTF silencing leads to decreased glycolytic gene expression, we infected K562 cells with a lentivirus encoding shRNAs against *MYC* or *MYB*. Silencing either *MYC* or *MYB* significantly decreased expression of a panel of glycolytic genes (Fig. S11D). These data demonstrate the control of glycolytic gene expression by LSTFs extends beyond cells of the melanocyte lineage.

To begin to test the hypothesis that this is a general phenomenon, we performed two complementary analyses. First, we used gene expression data from DepMap to determine the relationship between LSTFs and glycolytic gene expression. Because *MYC* is broadly expressed across these cancer cell lines and is a known transcriptional activator of glycolytic gene expression, apparent LSTF-glycolytic gene associations could be confounded by co-expression of *MYC* in the same cells. We therefore included *MYC* expression as a covariate in all regression models, allowing us to identify LSTF–glycolytic gene associations that are independent of *MYC*. Intra-lineage regression identified significant associations between LSTF expression and glycolytic genes for every LSTF tested, while also recapitulating examples of known biology, including the association of *KLF1* with *PKLR*, the erythroid-specific pyruvate kinase isoform (Fig. 4E, S12A). Even among LSTFs operating in the same lineage (*KLF1, SPI1, MYB* in hematopoietic cells), the identity and direction of significant associations differed, reflecting LSTF-specific rather than generic lineage-level regulation (Fig. 4E). This specificity extends to isoenzyme preferences, where *KLF1*- and *SPI1*-high cells favor distinct hexokinase and enolase isoforms (Fig. S12B). To ask whether LSTFs or *MYC* better account for the overall pattern of glycolytic gene expression, we summarized each lineage’s glycolytic expression profile using its first principal component (PC1), which captures the dominant axis of coordinated glycolytic variation, and measured the correlation between each LSTF and PC1 after statistically removing *MYC*’s contribution from both. In every lineage tested, this *MYC*-adjusted LSTF correlation with glycolytic PC1 exceeded the raw *MYC* correlation with PC1 in the same cells, demonstrating that LSTFs explain coordinated glycolytic gene expression independently of *MYC* and to a greater extent (Fig. 4F).

Second, we queried ChIP-Atlas across 32,000 distinct ChIP-Seq datasets for transcription factors that bind within 5kb of each metabolic gene transcriptional start site (TSS) (*42, 43*). To quantify the association between LSTF status and binding at metabolic gene TSSs, we calculated the adjusted odds ratio using a logistic regression model accounting for the total peak counts per experiment. LSTFs were significantly more likely to show enrichment at metabolic gene TSSs compared to non-LSTFs (Fig. 4G, S13A,B) in both human (adjusted OR = 3.10, 95% CI [2.86, 3.35], *P* = 5.80e-177) and mouse (adjusted OR = 2.61, 95% CI [2.20, 3.11], *P* = 9.28e-28). Taken together, these data suggest a generalized phenomenon wherein lineage-specifying transcription factors directly control metabolism through binding and controlling gene expression, thereby linking the regulation of specialized cellular function with the establishment of the requisite metabolic program.

## Discussion

Powerful methods such as CROP-seq (*44*) and Perturb-Seq (*45, 46*), have emerged for deciphering molecular phenotypes by combining genetic perturbation with scRNA-sequencing. These approaches can be used to decode gene regulatory mechanisms from top-down, defining the complex transcriptional response to one genetic perturbation. However, these experiments often require preselection of a relatively small number of genetic manipulations. Recently, more complete screens have been conducted, however the number of cells measured by droplet-based sequencing totals in the millions and the cost and computational demands can be prohibitive. As a result, these assays then do not easily lend themselves to gene by environment studies, where genetic perturbations are analyzed across environmental conditions, such as a small molecule, metabolic intervention, or infection. We demonstrate that TRoUT-FISH provides a bottom-up alternative that democratizes genetic screening for mediators of gene expression phenotypes and successfully deploy it across conditions of drug-induced transcriptional changes and unperturbed states.

Eukaryotic gene regulation is complex with multiple layers of control that impact the abundance of each transcript. We show that TRoUT-FISH effectively identifies transcriptional regulators, but as it uses RNA abundance as a molecular handle, TRoUT-FISH can also identify regulators that act outside of direct transcriptional control. Indeed, it should be able to discover factors that impact RNA abundance at any point in the RNA lifecycle, from signaling upstream of transcription to RNA degradation. As an example, our data suggest that *RCOR1, ZNF592*, and *LARP1* regulates glycolytic gene expression (Fig. S10B). *RCOR1* and *ZNF592* are interaction partners (*47*) and components of the transcriptional coregulator protein complex (*48*) which predominately control gene expression via histone modifications. LARP1 is an RNA-binding protein that post-transcriptionally regulates RNA stability and was recently shown to control metabolic genes and glycolysis (*49, 50*). Combined with the current work demonstrating direct control by transcription factors, we believe that TRoUT-FISH can identify regulators that act at any step to control mRNA abundance. In addition, while our current work focuses on mRNA, the method should be amenable to other RNA molecules including non-coding RNAs. In sum, TRoUT-FISH has the potential to provide deep data across many genetic perturbations for a set of target RNAs, with particular utility in identifying factors controlling transcription or transcript stability directly and decoding the mechanism of action for small molecules.

This study provides a compelling example of the intimate link between metabolism and cell fate (*36, 51, 52*), by showing that transcription factors that define cell fate and identity also directly drive expression of glycolytic genes. Our data suggest that this is essential for cellular function as inhibiting central carbon metabolism disrupts the specialized biochemistry of that lineage, thereby establishing a self-reinforcing relationship between the regulatory state and metabolic program of a cell (*52, 53*). This might suggest a model in which activation of SOX10 and MITF initiate a program for melanosome biogenesis and the enzymes responsible for production of pigment, while simultaneously activating the metabolic program to support growth, biomass, and energy required for these bespoke processes. Previous cancer studies have shown that the tissue of origin is a more important determinant of metabolism than the oncogenic driver (*54*) and that metastatic cells are metabolically more similar to their normal tissue of origin than to the tissue of residence (*55*). These studies demonstrate that some cell autonomous factors dictate the metabolic phenotype. Based on the results presented herein, we propose that this could be ascribed to the respective LSTFs, which we show to enforce a metabolic program alongside driving cellular identity. Genome-wide genetic screens in transformed cells have revealed that transcription factors, particularly those defining a given lineage, are frequently essential (*56, 57*). Hypotheses for this essentiality include oncogene addiction and the maintenance of a pro-growth cellular state. However, it is now tempting to speculate that their essentiality may in part reflect their role in driving essential metabolic functions (Fig. 4G).

It is evident that LSTFs do not bind or regulate all metabolic genes indiscriminately (Fig. 4E-G, S5C-D, S7E-H). Blood LSTFs bind enhancers to regulate hemoglobin and activate specific protein isoforms including PKLR; skin LSTFs control melanin biosynthetic genes, perhaps suggesting that a subset of specialized metabolic genes is privileged and selectively regulated by LSTFs. Other genes, particularly glycolytic genes (an ancient metabolic pathway), exhibit multi-factor regulation and are susceptible to tuning by many upstream factors. Our finding that *MYC*, a ubiquitous transcriptional activator broadly expressed across cancer cell lines, correlates with overall glycolytic gene expression yet is consistently outperformed by LSTFs (Fig. 4F) suggests a layered regulatory model: widely expressed factors such as *MYC* establish a basal level of glycolytic gene expression, while LSTFs fine-tune expression in response to the specific demands of the lineage (Fig. 4E-G, S12A-B). Our data reveal increased chromatin accessibility at glycolytic gene-adjacent regions containing AP1 motifs (Fig. 3F yellow), or RUNX and HAND motifs upon skin LSTF silencing (Fig. 3F, S6B-D purple). This might suggest a lineage reprogramming from a melanocyte-like state towards a neural crest-derivative state (*30, 58*–*60*); by eliminating the native lineage regulator, new factors are able to exert influence.

Specialization enables the diverse population of cells in the body to perform the wide array of unique functions that are required for the life of a complex organism. It has long been appreciated that specialization is controlled, in part, by lineage-specifying transcription factors that orchestrate transcriptional programs to drive the transition from progenitor to specialized cell types. These functions—such as novel organelle biogenesis or heme transport—are critical for organismal life and can require distinct metabolic adaptations, including melanin biosynthesis or mitochondria ejection. It follows that the same forces establishing the transcriptional programs that encode the structural proteins that execute these functions may also tune the cellular metabolic program to support them.

## Supporting information

DataS3

## Acknowledgements

We thank members of the Rutter lab for helpful discussions and comments on the manuscript. We thank members of the University of Utah Flow Cytometry core facility for their assistance. We thank members of the University of Utah High-Throughput Genomics core facility for their assistance. This work was supported by K00CA212445 to AJB and R35GM131854 to JR and the Huntsman Cancer Institute NC190504 to AJB and JR. JR is an investigator of the Howard Hughes Medical Institute. AJB and JR have filed a patent related to this work.

## Author contributions

Conceptualization: AJB, JR. Formal analysis: AJB, JYC, CNC, JGV. Funding acquisition: JR. Investigation: AJB, JYC, CNC, JGV, PW, AAC. Methodology: AJB. Project administration: JR, AJB. Resources: DEA, SPG, JR. Supervision: JR. Validation: AJB, JYC, CNC, JGV. Visualization: AJB. Writing – original draft: AJB, JR. Writing – review and editing: JR, AJB.

## Methods

### Oligonucleotide Sequences

All oligonucleotide sequences are provided in a supplementary document (Supplementary table 1). Oligo sequences include: 1) sgRNA sequences, 2) shRNA sequences, 3) qPCR primer sequences, 4) DNA oligos for RNA FISH.

### Plasmids and Libraries

The following plasmids were acquired from Addgene:

1. lentiGuide-Puro (*61*) was a gift from Feng Zhang (Addgene plasmid # 52963; http://n2t.net/addgene:52963; RRID:Addgene_52963)
2. lentiCas9-Blast (*61*) was a gift from Feng Zhang (Addgene plasmid # 52962; http://n2t.net/addgene:52962; RRID:Addgene_52962)
3. lenti_dCas9-KRAB-MeCP2 (*27*) was a gift from Andrea Califano (Addgene plasmid # 122205; http://n2t.net/addgene:122205; RRID:Addgene_122205)

The following CRISPR libraries were acquired from Addgene:

1. Lenti-human-TF-gRNA-puromycin library (*14*) was a gift from Chunliang Li (Addgene #162275; http://n2t.net/addgene:162275; RRID:Addgene_162275)
2. Human Epigenetic Knockout Library (*17*) was a gift from Kivanc Birsoy (Addgene #162256; http://n2t.net/addgene:162256; RRID:Addgene_162256)

### Antibodies

Antibodies against p21 (Catalog: #2947) and p53 (Catalog: #2524) were purchased from Cell Signaling Technology. Antibodies against PMEL were purchased from ThermoFisher (Catalog: MA1-34759). Antibodies against MITF for ChIP-sequencing were purchased from Active Motif (Catalog: 39789, RRID: AB_2614955).

### Cloning

CRISPRi sgRNAs were selected from previously designed whole genome sets (*26*) or designed using CRISPick (Broad Institute). CRISPR nuclease sgRNAs were designed using CRISPOR (*62*) for selected exons. Sequences were appended with flanking sequences for cloning into BsmBI-digested lentiGuide-Puro or lentiCRISPRv2. In brief, reverse complement oligonucleotides with appropriate flanking sequences were resuspended in water. Oligo pairs were phosphorylated with T4 Polynucleotide Kinase then annealed in a thermocycler. Oligos were incubated with T4 PNK at 37C for 30 minutes, then heated to 95C for 5 minutes, with a 5C step down in temperature with a 1-minute hold until reaching 25C. Oligos were then diluted 200-fold and ligated into predigested and dephosphorylated lentiGuide-Puro or lentiCRISPRv2 with T4 DNA ligase.

### Cell Culture

HEK293T, HCT116, and K562 cells were obtained from ATCC. MeWo cells were a gift from Dr. Martin McMahon. B16F10 cells were a gift from Dr. Minna Roh-Johnson. HCT116, K562, and MeWo cells were cultured in RPMI supplemented with 10% fetal bovine serum, 100 units/ml penicillin, and 100 µg/ml streptomycin. HEK293T and B16F10 cells were cultured in DMEM supplemented with 10% fetal bovine serum, 100 units/ml penicillin, and 100 µg/ml streptomycin. HEK293T, HCT116, and MeWo cells were selected with 1 µg/mL puromycin or 10 µg/mL blasticidin. K562 cells were selected with 2.5 µg/mL puromycin or 20 µg/mL blasticidin.

For ^13^C-glucose tracing experiment, MeWo cells were cultured in RPMI media without glucose with 10% in-house dialyzed fetal bovine serum, 100 units/ml penicillin, and 100 µg/ml streptomycin. Briefly, 250 mL of serum was aliquoted into 3.5kDa SnakeSkin Dialysis Tubing and dialyzed against 20 volumes of PBS. PBS was changed every 12 hours for a total dialysis of 48 hours after which serum was aliquoted and frozen until use.

### Virus Production and Infection

Lentiviruses were produced in HEK293T cells by transfecting plasmids and packaging plasmids (psPAX2 and VSVG) with polyethylenimine (PEI) transfection reagent. Media was collected every 24 hours for 96 hours total. Viral supernatant from each timepoint was combined and syringe filtered with 0.45 µm PES filters and stored at 4C until use. Filtered viral supernatant was applied to target cells at a ratio of 2:1 (viral media:fresh media) with 10 µg of polybrene.

### Immunoblotting

Cell lysates were prepared in 1% sodium deoxycholate, 0.1% SDS, 1% Triton X-100, 0.01 M Tris pH8.0, 140 mM NaCl. Protein lysates were separated by SDS-PAGE and transferred to nitrocellulose membranes by wet transfer. All primary antibodies were incubated overnight at 4C. Alexafluor-conjugated goat anti-rabbit (IRDye800, Rockland) or goat anti-mouse (Alexafluor-680, Life Technologies) antibodies were used as secondary antibodies (1:10,000). Immunoblots were developed using an Odyssey Imager (LI-COR).

### Metabolomics

Cells were plated in 100 mm dishes, washed three times with PBS, and cultured with media containing [U-^13^C]glucose for 2 hours. After incubation, medium was collected for analysis, and cells were washed twice with ice-cold 0.9% saline solution. Pre-chilled extraction solvent (80% methanol in water at −80C) was added to each plate and plates were transferred to −80C for 15 minutes. While kept on dry ice, cells were then scraped into the cold solvent. Extracts were centrifuged at 20,000xg at 4C for 10 min, and the supernatant transferred to a new 1.5 mL tube. Solvent in was removed by SpeedVac, and dried extracts stored at −80C until LC-MS analysis.

Polar extracts were separated by HILIC on a Vanquish HPLC (ThermoFisher) with an XBridge BEH Amide column (2.1 × 150 mm, 2.5 µm, Waters). Mobile phase A was 20 mM NH_4_OH + 20 mM NH_4_OAc in 95:5 acetonitrile:water (pH 9.5); B was 100% acetonitrile. The flow rate was 150 µL/min, and the gradient proceeded as follows: start at 90% B for 2 min, drop to 75% B at 3 min (hold to 7 min), to 70% at 8 min (hold to 9 min), to 50% at 10 min (hold to 12 min), to 25% at 13 min (hold to 14 min), to 0% at 16 min (hold to 20.5 min), then return to 90% B by 21 min and re-equilibrate to 25 min. The autosampler was

4 °C, the column 30 °C, and injection volume 3 µL. Detection was by negative-mode electrospray on a Q Exactive HF Orbitrap (75,000 resolution at m/z 200; scan range 75–1000 m/z). Data were processed with EL-MAVEN, peaks assigned by exact mass against known standards (*63*), and intensities corrected for natural isotoptic abundance using the R package AccuCor (*64*).

### IncuCyte

B16F10 cells were seeded in 48-well plates and allowed to adhere overnight before treatment with either vehicle or UK5099 (10 µM). Phase contrast images were acquired every 8 hours for 4 days. Confluence was determined using Satorious Incucyte software.

### RNA Extraction, cDNA Synthesis, and Quantitative Polymerase Chain Reaction

RNA was extracted from 6cm plates with 300µL of TRIzol solution (ThermoFisher, Catalog: 15596018). RNA was isolated using the Direct-zol RNA Miniprep kit with on-column DNase treatment (Zymo Research, Catalog: R2052). 2µg of RNA were used for complementary DNA (cDNA) synthesis using the High-Capacity cDNA Reverse Transcription Kit (ThermoFisher, Catalog: 4368813). cDNA was diluted 10-fold before being quantified via quantitative PCR (qPCR) using the QuantStudio7 Flex system with Power SYBR Green Master Mix (ThermoFisher, Catalog: 4368708). Experiments were performed in biological and technical triplicate, and the genes POLR2A and RPLP0 were used as reference genes during analysis. Data were analyzed using the comparative C_T_ (ΔΔCt) method.

### Melanin Assay

Cell pellets were thawed on ice and resuspended in 100µL 1N NaOH + 10%DMSO. Resuspended pellets were vortexed briefly and then incubated at 80C for 90 minutes with constant rotation to extract cellular melanin. Cell debris was pelleted by centrifugation at 18000xg for 10 minutes, after which the supernatant was transferred to a fresh tube. A standard curve was set up using synthetic melanin (range 1000µg/mL to 7.8µg/mL). Synthetic melanin was purchased from Sigma (Catalog: M8621). 50µL of each standard was plated on a 96-well plate. Samples were plated in technical duplicate (50µL per well). Absorbance was read at 490nm.

### SABER-FISH PER

Primer Exchange Reaction (PER) was carried out as described previously (*11–13*). In brief, oligonucleotides were designed according as described previously (*65, 66*) to ensure transcriptome wide specificity. DNA oligonucleotides targeting RNAs of interest were appended with single-stranded DNA tails via catalytic PER hairpins. Sequences for each PER are listed in a supplementary table (Supplementary table 1).

### TRoUT-FISH screens

Lentiviruses were produced in HEK293T cells by transfecting plasmids and packaging plasmids (psPAX2 and VSVG) with polyethylenimine transfection reagent. Media was collected every 24 hours for 96 hours total. Media was pooled, filtered with 0.45 µm PES filters, aliquoted, and frozen at −80C. The titer of lentivirus supernatant was determined by infecting target cells at various amounts of virus with polybrene followed by spinfection (30C for 60 minutes at 1200 xg). The next day cells were reseeded and treated with puromycin. Viability was assessed 3 days post infection. Cells were infected at a Multiplicity of Infection (MOI) of ~0.4 via spinfection, and treated with puromycin 24 hours post infection and puromycin selection persisted for 7 days. Depending on the experiment, cells were either treated with Nutlin-3a for 6 hours followed by fixation or given fresh media for 6 hours followed by fixation with cold 4% PFA.

Cells were permeabilized with PBS containing 0.1% Triton X-100 and Hoechst 33342 for 10 minutes rocking at 4C. Cells were incubated with 43C wHyb buffer (*12, 13*) for 30 minutes, then incubated overnight at 43C in hybridization buffer containing SABER-FISH PER purified DNA oligonucleotides. Subsequently, cells were washed two times with 43C wHyb buffer, one time with 2X SSC buffer, and incubated at 30C for 1 hour in branch hybridization buffer containing SABER-FISH PER purified branch DNA oligonucleotides. After branching, cells were washed with 2X SSC and 37C PBS before incubation for 10 minutes in PBS containing fluorescently labeled secondary DNA oligonucleotides. After staining, cells were washed two times with 37C PBS and resuspended in 0.1% BSA (Millipore, Catalog: 126609-5GM) in PBS for flow cytometry. Events were gated based on FSC and SSC to remove debris, then gated on DNA dye (violet laser) to ensure only singlets were sorted. A reference gene (EEF2) was used as a control to ensure cell quality and adequate staining. Cells were sorted based on fractions of the target gene distribution, sorting the lowest 10% and highest 10% of the total population. An unsorted population was also acquired. After sorting cells were incubated in genomic DNA isolation buffer (0.5% SDS, 10 mM TRIS pH 7.5, 10 mg/ml Proteinase K) and incubated at 56C for ≥ 36 hours with gentle agitation. Genomic isolation was performed with the Qiagen DNA Mini Kit (Catalog: 51304). Libraries were generated by PCR amplification and sequenced by 150bp paired-end sequencing at the University of Utah High-Throughput Genomics Core Facility. Sequencing reads were quantified using BBtools Seal and analysis was performed by MAGeCK (*67, 68*).

### ChIP-Sequencing

ChIP-sequencing was performed as previously described with slight modifications (*69*). In brief, MeWo cells were seeded in a 15cm plate at 1e7 cells per plate. 6 hours prior to fixation, media was replaced with 15mL of fresh RPMI media. Cells were fixed at a final concentration of 1% PFA for 5 minutes. Fixation was quenched with 0.125M final concentration of glycine for 10 minutes at RT. Media was aspirated, and then cells were washed twice with cold PBS containing 1 mM PMSF. Cells were scrapped, pelleted, flash frozen, and stored at −80C until use. Chromatin was sonicated in a Bioruptor Pico sonicator for 30 minutes according to the following regimen: 10 cycles of 30 seconds on and 30 seconds off were followed by a 10-minute rest period, repeated 3 times. 5µg of Active Motif MITF Ab was added to sonicated chromatin and incubated overnight at 4C. 10% of chromatin was reserved as input DNA. 30µL of magnetic Dynabeads (Protein G) were used for each IP and rotated for 4 hours at 4C. Magnetic beads were washed and chromatin was eluted followed by incubation at 65C for 16 hours to reverse crosslinks and subquently treated with RNAse A and Proteinase K. DNA was purified by Phenol-Chloroform extraction and GlycoBlue Co-precipitate (ThermoFisher, Catalog: AM9515). DNA was then cleaned and concentrated using the Zymo Clean and Concentrate kit (Zymo Research, Catalog: D4003). Libraries were generated by NEBNext Ultra II DNA Library Prep Kit. Libraries were sequenced by 150bp paired-end sequencing at the University of Utah High-Throughput Genomics Core Facility. Data analysis used the ENCODE ChIP-sequencing pipeline (*70*) and visualized with IGV.

### RNA-Sequencing

RNA was isolated as described above. Libraries were generated by NEBNext Ultra II Directional RNA Library Prep with rRNA Depletion Kit or Illumina TruSeq Stranded mRNA Library Preparation Kit with polyA selection.

Libraries were sequenced by 150bp paired-end sequencing at the University of Utah High-Throughput Genomics Core Facility. Reads were trimmed of adaptor sequences using Trim Galore (*71*) and aligned to hg38 with STAR (*72*) and differentially expressed genes were identified using DESeq2 (*73*) version 1.30.0.

### ATAC-Sequencing

MeWo_KRAB cells were transduced with control sgRNAs or those designed to silence MITF or SOX10 and selected with puromycin 24 hours later. Four days post infection, cells were prepped for ATAC-sequencing as previously described (*74*). In brief, nuclei were isolated from target cells and genomic DNA underwent transposition using Tagment DNA Enzyme 1 (Illumina Catalog: 15027865). Sequencing libraries were generated by PCR amplification with cycles measured by qPCR to avoid saturation. Libraries were purified by double-sided AMPure XP bead purification (Beckman Coulter, Catalog: A63880). Libraries were sequenced by 150bp paired-end sequencing at the University of Utah High-Throughput Genomics Core Facility. Data analysis used the ENCODE ATAC-sequencing pipeline (*70*) and visualized with IGV or deepTools (*75*) for heatmaps. Differentially accessible regions were identified by extracting and counting all genomic intervals using featureCounts (*76*) and DESeq2. Motifs from differentially accessible regions were identified from using HOMER (*32*).

### DepMap lineage analysis

DepMap Public 26Q1 release was used for gene expression (log_2_(TPM+1)), CRISPR dependency (Chronos scores), copy number, and cell line annotations. Cell lines with complete data were used (n = 1,140) and lineage-specifying transcription factors were selected based on known roles (SOX10 and MITF (skin/melanoma), KLF1, SPI1, and MYB (hematopoietic), HNF1B (kidney)). MYC was included in parallel as a comparison. For group-based analyses, a cell line was only considered part of the LSTF group if: LSTF expression was ≥ 75^th^ percentile expression across all 1,140 cell lines, and a gene dependency score < −0.5. If a cell line could be assigned to more than one LSTF it was excluded.

To perform intra-lineage regression, each LSTF was fit with ordinary least-squares for each of the 56 metabolic genes (Supplementary table 2) across all cell lines in that lineage with MYC included as a covariate. The beta coefficient and standard error for the LSTF term are reported and multiple-testing correction using BH was applied with a significance threshold of FDR < 0.1. For HNF1B (n = 32 kidney lines), leave-one-out analysis was used to confirm sign stability. To assess the contribution of LSTF vs MYC on glycolytic gene expression as a whole, the glycolytic gene expression was extracted for all cells within a lineage and z-scored using prcomp in R and the first principal component (PC1) was used. Two linear models were fit: (1) PC1 ~ MYC and (2) LSTF ~ MYC. The partial correlation between LSTF and PC1, while controlling for MYC, was computed by correlating the residuals of model and (2). The absolute value of the partial correlation is reported to remove sign ambiguity inherent to PCA orientation. Claude Sonnet 4.6 was used for some part of this analysis.

### ChIP-Atlas analysis

The ChIP-Atlas database was queried using the enrichment analysis module. Three gene sets were used for each human and mouse including metabolic genes (*77*), genes related to positive regulation of transcription by RNA polymerase II (GO: 0045944), or random genes matching the set size of the metabolic gene list. For each gene set, logistic regression was used to estimate the odds of a ChIP-sequencing experiment showing fold enrichment greater than 1 (i.e., greater overlap with the target gene set than expected by chance) was modeled as a binary outcome for LSTF or non-LSTF experiments with total peak count for each study included as a covariate. The adjusted odds ratio and confidence intervals were estimated from the exponentiated model coefficients. Logistic regression was performed in R (4.2.2) using the glm() function with a binomial family. Odds ratio and 95% confidence intervals were obtained using the tidy() function from the broom package (1.0.3). Transcription factors were classified as LSTFs based on established roles in lineage specification, requiring both transcription factor identity and tissue context in the ChIP-Atlas data (Supplementary table). Random gene sets were generated by sampling 3000 genes from the appropriate species coding gene set, matched to the size of the metabolic gene list.

### Sample Preparation for Mass Spectrometry

Samples for protein analysis were prepared essentially as previously described (*78, 79*). Proteomes were extracted using a buffer containing 200 mM EPPS pH 8.5, 8M urea, 0.1% SDS and protease inhibitors. Following lysis, 150 µg of each proteome was reduced with 5 mM TCEP. Cysteine residues were alkylated using 10 mM iodoacetamide for 20 minutes at RT in the dark. Excess iodoacetimide was quenched with 10 mM DTT. A buffer exchange was carried out using a modified SP3 protocol (*80*). Briefly, ~1500 µg of Cytiva SpeedBead Magnetic Carboxylate Modified Particles (65152105050250 and 4515210505250), mixed at a 1:1 ratio, were added to each sample. 100% ethanol was added to each sample to achieve a final ethanol concentration of at least 50%. Samples were incubated with gentle shaking for 15 minutes. Samples were washed three times with 80% ethanol. Protein was eluted from SP3 beads using 200 mM EPPS pH 8.5 containing Lys-C (Wako, 129-02541). Samples were digested overnight at room temperature with vigorous shaking. The next morning trypsin (ThermoFisher Scientific) was added to each sample and further incubated for 6 hours at 37º C. Following digestion, an equal volume of each sample was pooled to generate a pooled sample to be used as a bridge between two TMTPro 16-plex experiments. Acetonitrile was added to each sample to achieve a final concentration of ~33%. Each sample was labelled, in the presence of SP3 beads, with ~300 µg of TMTPro reagents (ThermoFisher Scientific). Following confirmation of satisfactory labelling (>97%), excess TMT was quenched by addition of hydroxylamine to a final concentration of 0.3%. The full volume from each sample was pooled and acetonitrile was removed by vacuum centrifugation for 1 hour. The pooled sample was acidified and peptides were de-salted using a Sep-Pak 50mg tC18 cartridge (Waters). Peptides were eluted in 70% acetonitrile, 1% formic acid and dried by vacuum centrifugation.

### Basic pH reversed-phase separation (BPRP)

TMT labeled peptides were solubilized in 5% ACN/10 mM ammonium bicarbonate, pH 8.0 and ~300 µg of TMT labeled peptides were separated by an Agilent 300 Extend C18 column (3.5 μm particles, 4.6 mm ID and 250 mm in length). An Agilent 1260 binary pump coupled with a photodiode array (PDA) detector (Thermo Scientific) was used to separate the peptides. A 45-minute linear gradient from 10% to 40% acetonitrile in 10 mM ammonium bicarbonate pH 8.0 (flow rate of 0.6 mL/min) separated the peptide mixtures into a total of 96 fractions (36 seconds). A total of 96 Fractions were consolidated into 24 samples in a checkerboard fashion and vacuum dried to completion. Each sample was desalted via Stage Tips and re-dissolved in 5% FA/ 5% ACN for LC-MS3 analysis.

### Liquid chromatography separation and tandem mass spectrometry (LC-MS3)

Proteome data were collected on an Orbitrap Fusion Lumos mass spectrometer (ThermoFisher Scientific) coupled to a Proxeon EASY-nLC 1000 LC pump (ThermoFisher Scientific). Fractionated peptides were separated using a 120 min gradient at 550 nL/min on a 35 cm column (i.d. 100 μm, Accucore, 2.6 μm, 150 Å) packed in-house. MS1 data were collected in the Orbitrap (60,000 resolution; maximum injection time 50 ms; AGC 4 × 10^5^). Charge states between 2 and 6 were required for MS2 analysis, and a 120 s dynamic exclusion window was used. Top 10 MS2 scans were performed in the ion trap with CID fragmentation (isolation window 0.5 Da; Rapid; NCE 35%; maximum injection time 50 ms; AGC 2 × 10^4^). An on-line real-time search algorithm (Orbiter) was used to trigger MS3 scans for quantification (*81*). MS3 scans were collected in the Orbitrap using a resolution of 50,000, NCE of 55%, maximum injection time of 200 ms, and AGC of 3.0 × 10^5^. The close out was set at two peptides per protein per fraction (*81*).

### Proteomic data analysis

Raw files were converted to mzXML, and monoisotopic peaks were re-assigned using Monocle (*82*). Searches were performed using the Comet search algorithm against a mouse database downloaded from Uniprot in May 2021. We used a 50 ppm precursor ion tolerance, 1.0005 fragment ion tolerance, and 0.4 fragment bin offset for MS2 scans collected in the ion trap. TMTpro on lysine residues and peptide N-termini (+304.2071 Da) and carbamidomethylation of cysteine residues (+57.0215 Da) were set as static modifications, while oxidation of methionine residues (+15.9949 Da) was set as a variable modification. Each run was filtered separately to 1% False Discovery Rate (FDR) on the peptide-spectrum match (PSM) level. Then proteins were filtered to the target 1% FDR level across the entire combined data set. For reporter ion quantification, a 0.003 Da window around the theoretical m/z of each reporter ion was scanned, and the most intense m/z was used. Reporter ion intensities were adjusted to correct for isotopic impurities of the different TMTpro reagents according to manufacturer specifications. Peptides were filtered to include only those with a summed signal-to-noise (SN) ≥ 160 across all TMT channels. The signal-to-noise (S/N) measurements of peptides assigned to each protein were summed (for a given protein). These values were normalized so that the sum of the signal for all proteins in each channel was equivalent thereby accounting for equal protein loading.

### Electron Microscopy

B16F10 cells were seeded and treated with DMSO or UK-5099 for 48hrs. Once the cells reached confluence, the growth media was replaced with cold with fixative solution (2.5% glutaraldehyde, 1% PFA; 0.1 M Cacodylate buffer, pH 7.2). The cells were collected and incubated at 4°C for overnight. The next day, aclar disks with cells were gently rinsed twice with cacodylate buffer and post-fixed with a buffered 2% osmium tetroxyde for 1 hr at room temperature. Then they were rinsed with dH2O and pre-stained for 1 hr with saturated filtered uranyl acetate at room temperature. Dehydration and infiltration was performed using ethanol - graded series for 7 min each (50%; 70%; 95%-2x; 100%-3x) and pure acetone 3×7 min each. The cell pellet was infiltrated with 50% epon resin:acetone for 1 hr, then with 75% epon resin for overnight and 100% epon resin over 8 hrs with 3 changes. Cells were embedded and polymerized at 60°C for 48 hrs. Ultrathin (70nm) sections were obtained with diamond knife (Diatome) and an ultratome Leica UC6 (Leica Microsystems, Vienna, Austria). Grids with sections were post stained with saturated uranyl acetate in dH2O for 20 min and with lead citrate for 10 min and imaged at 120kV using JEOL-JEM 1400 Plus electron microscope (Tokyo, Japan).

---

**Figure S1:**
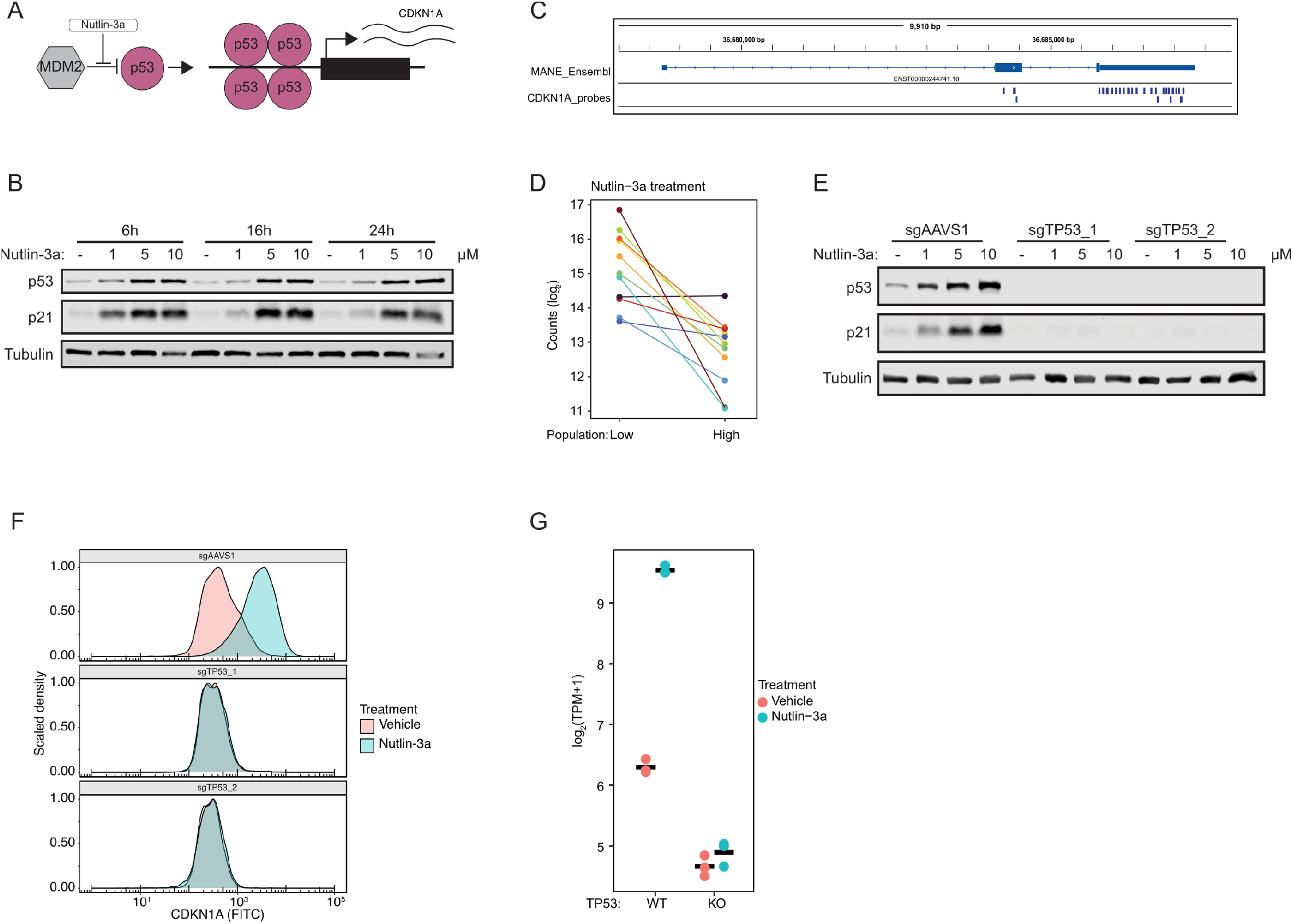
*TP53* controls *CDKN1A*. **(A)** Schematic of Nutlin-3a mechanism of action. **(B)** Genome browser visualizing binding sites of DNA oligonucleotides hybridizing to CDKN1A RNA. **(C)** Immunoblots of HCT116 cells treated with Nutlin-3a at the indicated doses and timepoint. **(D)** Dot plot showing TRoUT-FISH sgRNA counts against TP53 for CDKN1A RNA low and high populations in HCT116 cells treated with Nutlin-3a. Each colored point indicates an sgRNA. **(E)** Immunoblots of HCT116 cells treated with Nutlin-3a at the indicated doses for the indicated genotypes. **(F)** Density plot showing CDKN1A levels as measured by RNA FISH and flow cytometry. **(G)** Dot plot showing RNA-sequencing transcripts-per-million (GSE137297) of CDKN1A from WT or TP53-/-HCT116 cells treated with 5 µM Nutlin-3a for 6 hours.

**Figure S2:**
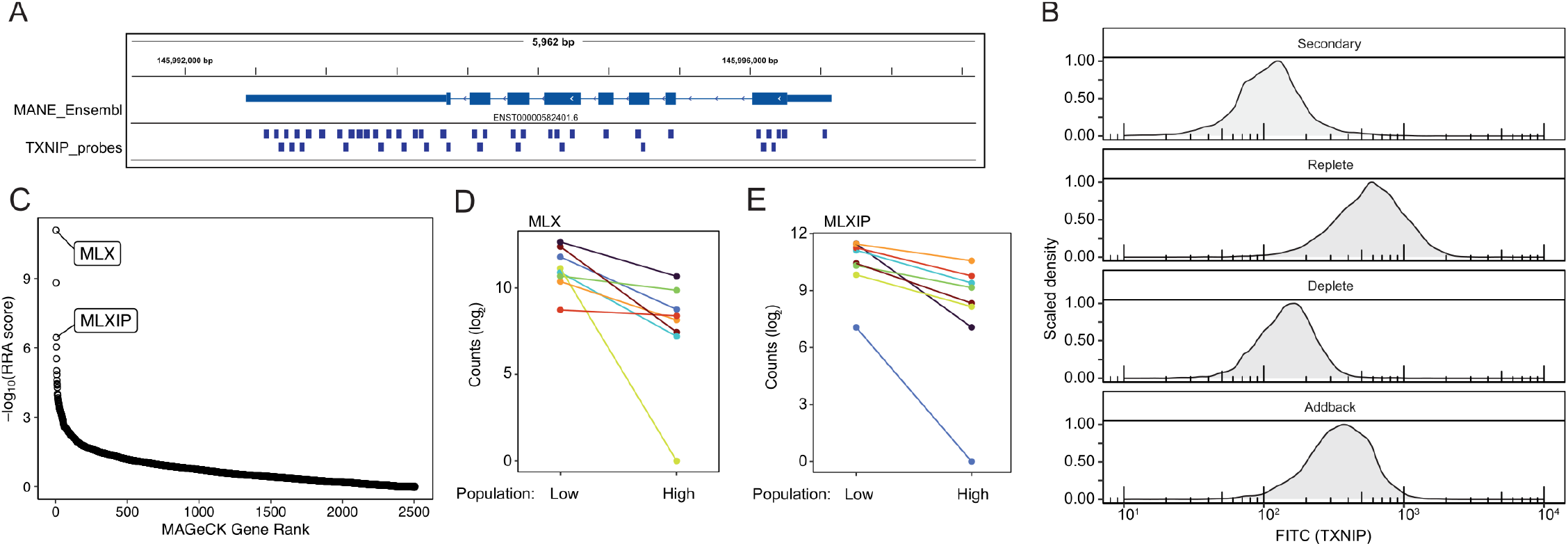
TRoUT-FISH identifies known regulators of *TXNIP*. **(A)** Genome browser visualizing binding sites of DNA oligonucleotides hybridizing to TXNIP RNA. **(B)** Density plot showing *TXNIP* levels as measured by RNA FISH and flow cytometry K562 cells cultured in the presence or absence of glucose for 16 hours with glucose addback. **(C)** TRoUT-FISH results for basal *TXNIP* abundance in K562 cells. **(D)** Dot plot showing TRoUT-FISH sgRNA counts against *MLX* for *TXNIP* RNA low and high populations in K562 cells as in **(C)**. Each colored point indicates an sgRNA. **(E)** Dot plot showing TRoUT-FISH sgRNA counts against *MLXIP* for *TXNIP* RNA low and high populations in K562 cells as in **(C)**. Each colored point indicates an sgRNA.

**Figure S3:**
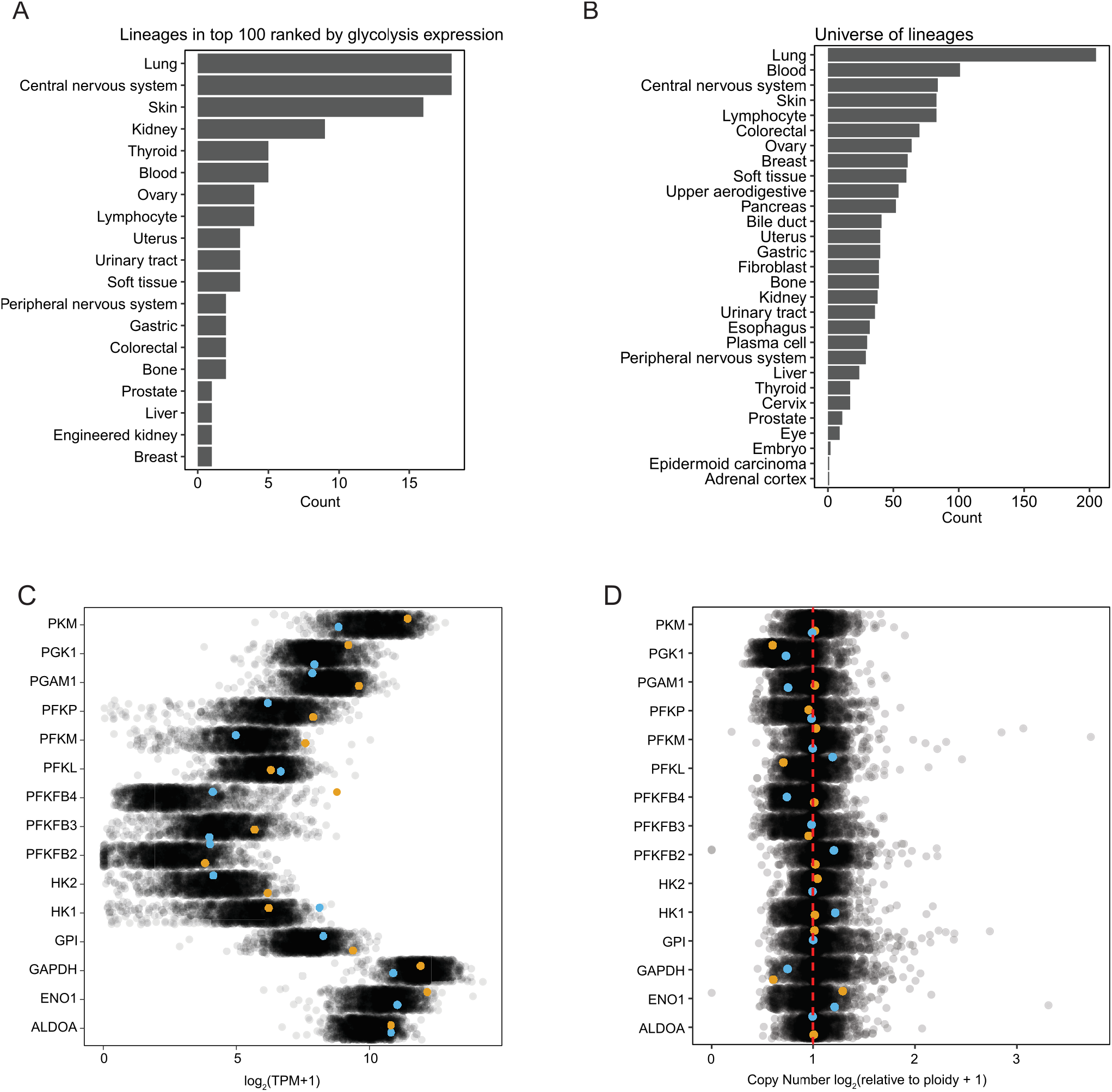
Profiling of DepMap gene expression reveals lineage enrichments. **(A)** DepMap transcript abundance data (21Q3) was analyzed. For each glycolytic gene, cells were ranked based on log_2_(TPM+1) abundance and assigned a rank. This was repeated for each gene. Cell lines lower cumulative scores (indicating higher expression) were analyzed for enrichment stratifying the top 100 cell lines. Bar plot showing the aggregate number of cell lines in the top 100 based on expression for each lineage or **(B)** bar plot showing the universe of cell lineages in the dataset. Fisher’s exact test was performed for skin (*P* = 1.832 × 10^−4^) and central nervous system (*P* = 2.1 × 10^−6^). **(C)** Scatterplot showing transcript abundance or **(D)** copy number, plotted for each cell line for each gene. Orange indicates MeWo and blue indicates K562 cells.

**Figure S4:**
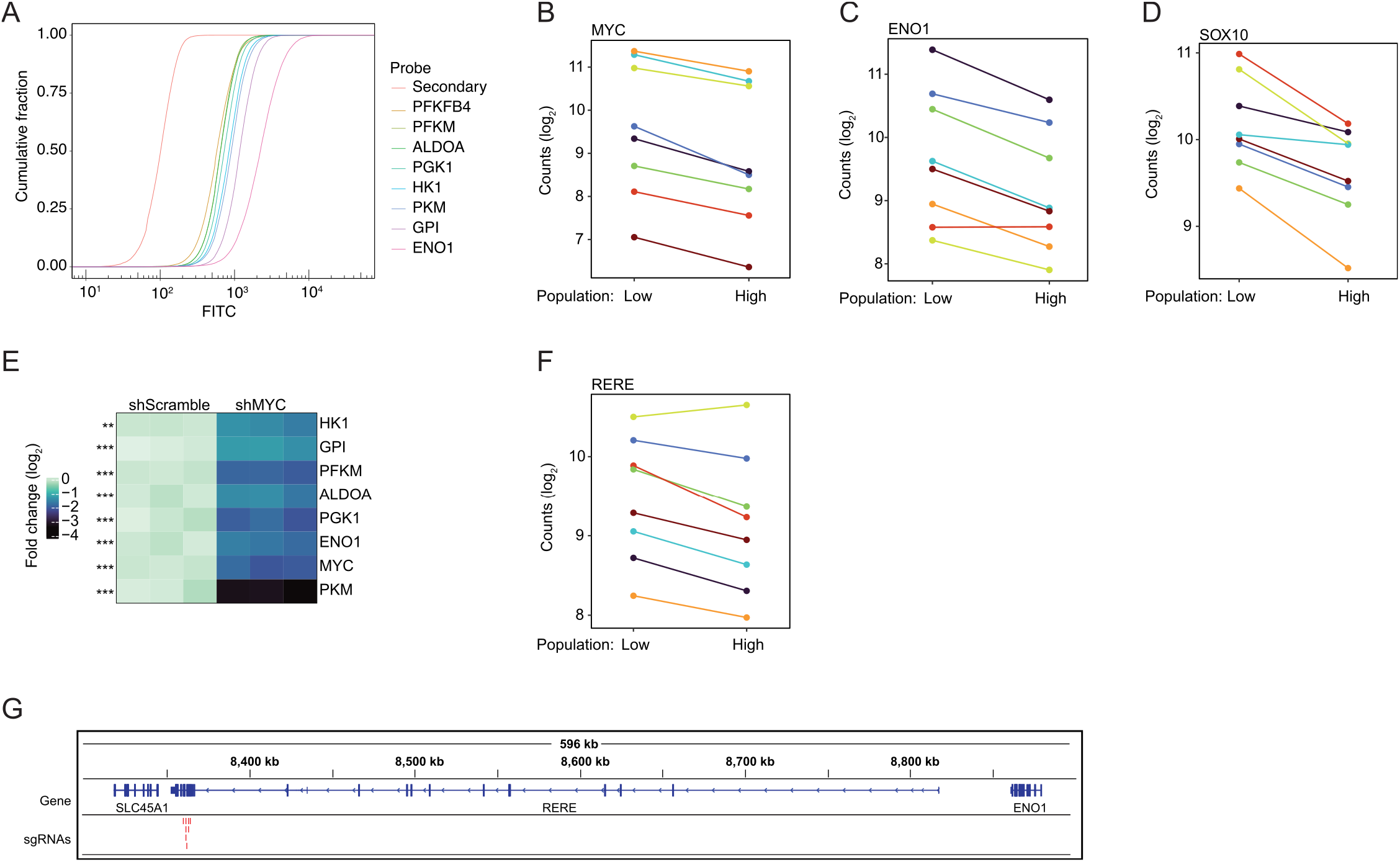
TRoUT-FISH identifies regulators of glycolytic gene expression. **(A)** Cumulative distribution plot showing RNA FISH signal for each indicated target gene in MeWo cells. **(B)** Dot plot showing TRoUT-FISH sgRNA counts against MYC for ENO1 RNA low and high populations in MeWo cells. Each colored point indicates an sgRNA. **(C)** Dot plot showing TRoUT-FISH sgRNA counts against ENO1 for ENO1 RNA low and high populations in MeWo cells. Each colored point indicates an sgRNA. **(D)** Dot plot showing TRoUT-FISH sgRNA counts against SOX10 for ENO1 RNA low and high populations in MeWo cells. Each colored point indicates an sgRNA. **(E)** Heatmap depicting the log_2_ fold change of the indicated target gene in MYC shRNA transduced MeWo cells as measured by qPCR. Experiments performed in triplicate. A two-tailed, two-sample t-test was performed (\**P* < 0.05, \*\**P* < 0.01, \*\*\**P* < 0.001). **(F)** Dot plot showing TRoUT-FISH sgRNA counts against RERE for ENO1 RNA low and high populations in MeWo cells. Each colored point indicates an sgRNA. **(G)** Genome browser tracks visualizing binding sites of DNA oligonucleotides hybridizing to RERE RNA.

**Figure S5:**
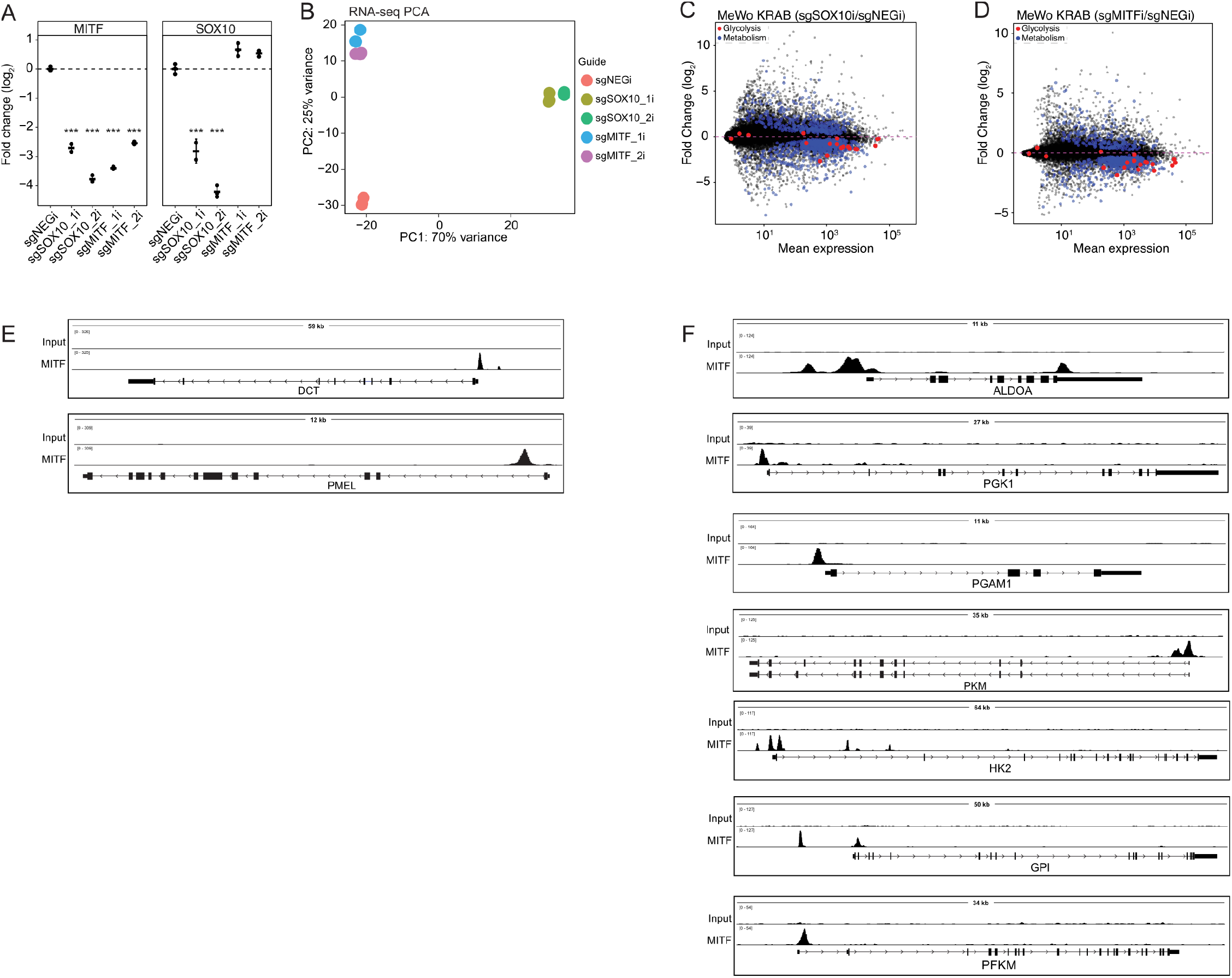
Silencing *SOX10* and *MITF* impacts metabolic gene expression. **(A)** Dot plot showing changes in MITF or SOX10 expression in MeWo_KRAB cells infected with guides against the indicated genes 4 days post infection, as determined by qPCR. A one-way ANOVA and Dunnett test were performed (\**P* < 0.05, \*\**P* < 0.01, \*\*\**P* < 0.001). **(B)** PCA plot from RNA-sequencing data as in **(A). (C)** MA-plot of MeWo KRAB sgSOX10i as in **(B)**. Glycolytic genes are highlighted in red and metabolic genes (FDR < 0.05) are highlighted in blue. **(D)** MA-plot of MeWo KRAB sgMITFi as in **(B)**. Glycolytic genes are highlighted in red and metabolic genes (FDR < 0.05) are highlighted in blue. **(E)** Genome browser tracks visualizing of ChIP-sequencing peaks of MITF binding in MeWo cells at canonical MITF target genes or **(F)** at glycolytic genes.

**Figure S6:**
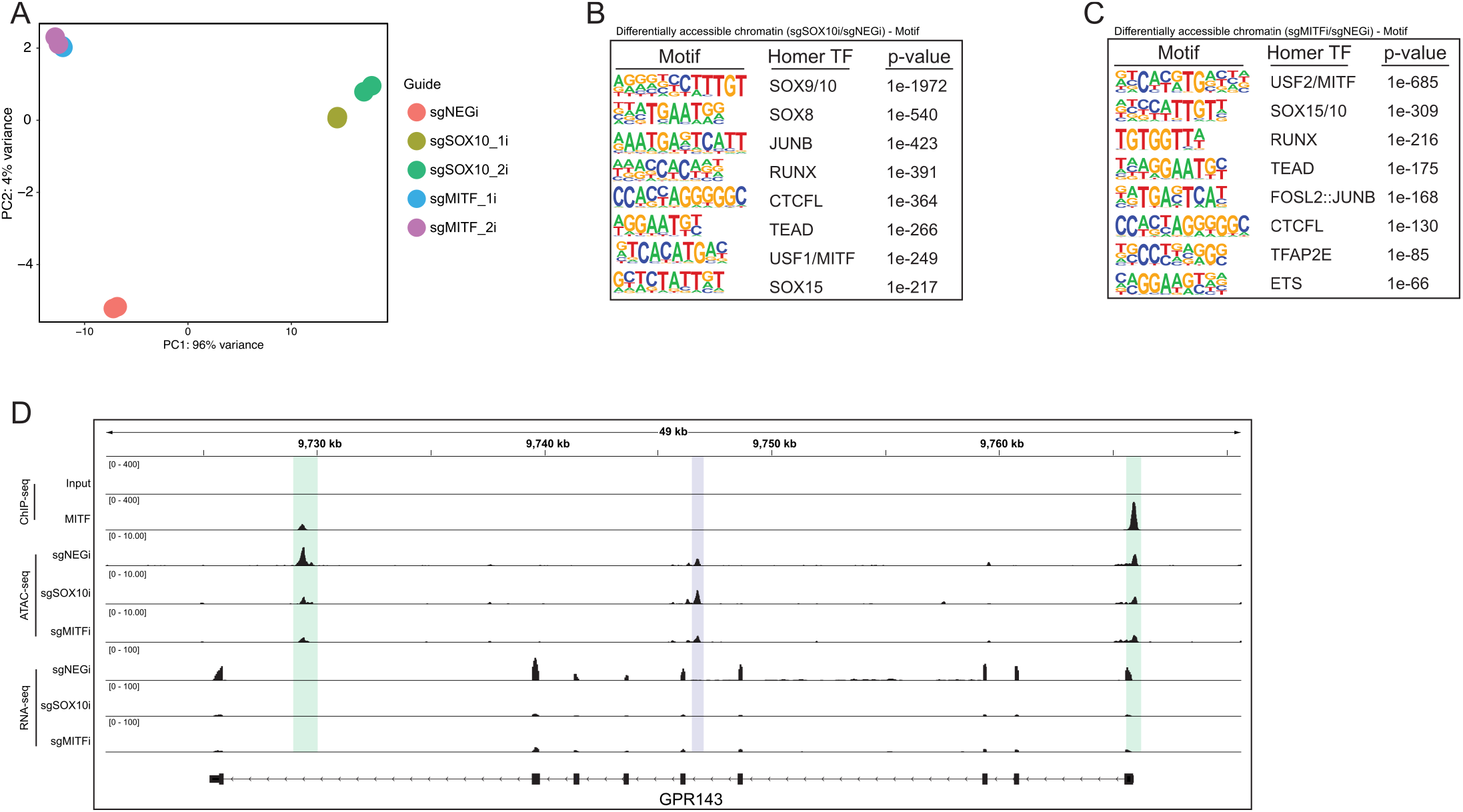
Silencing *SOX10* and *MITF* alters chromatin accessibility. **(A)** PCA plot from ATAC-sequencing of MeWo_KRAB cells transduced with guides against SOX10 or MITF 4 days post infection. **(B)** HOMER motif enrichment of differentially accessible chromatin of MeWo_KRAB sgSOX10i as in **(A). (C)** HOMER motif enrichment of differentially accessible chromatin of MeWo_KRAB sgMITFi as in **(A). (D)** Genome browser tracks visualizing MITF ChIP-sequencing, ATAC-sequencing, and RNA-sequencing in MeWo_KRAB cells. Purple region is increased accessibility with a RUNX motif and green region is decreased accessibility for either or both genotypes.

**Figure S7:**
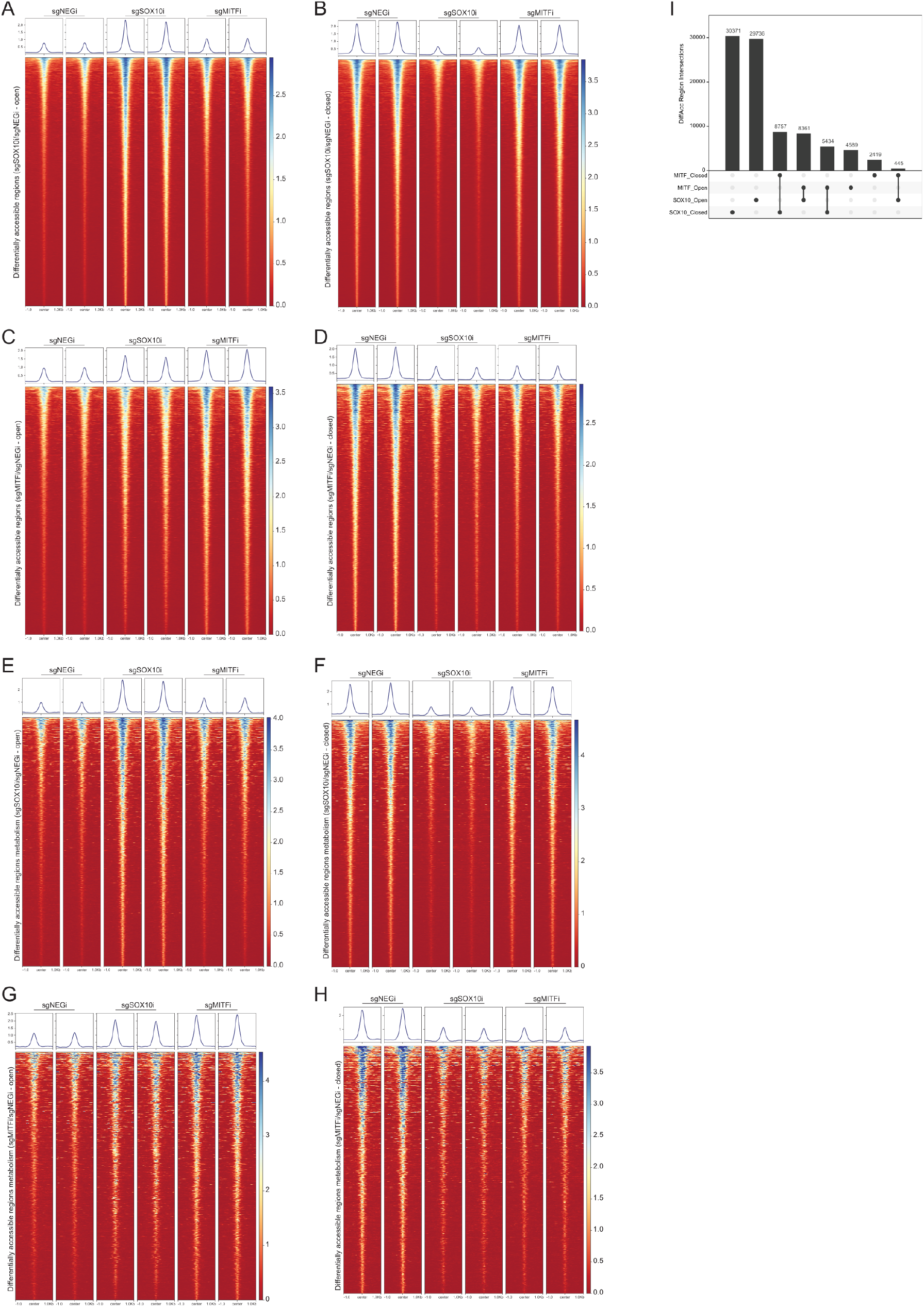
Silencing *SOX10* and *MITF* profoundly alter chromatin accessibility proximal to metabolic genes. **(A, B)** Density heatmap of peaks differentially open **(A)** or closed **(B)** after *SOX10* suppression. **(C, D)** Density heatmap of peaks differentially open **(C)** or closed **(D)** after *MITF* suppression. **(E, F)** Density heatmap of peaks associated with metabolic genes differentially open **(E)** or closed **(F)** after *SOX10* suppression. **(G, H)** Density heatmap of peaks associated with metabolic genes differentially open **(G)** or closed **(H)** after *MITF* suppression. **(I)** Upset plot representing overlap between differentially accessible regions between genotypes.

**Figure S8:**
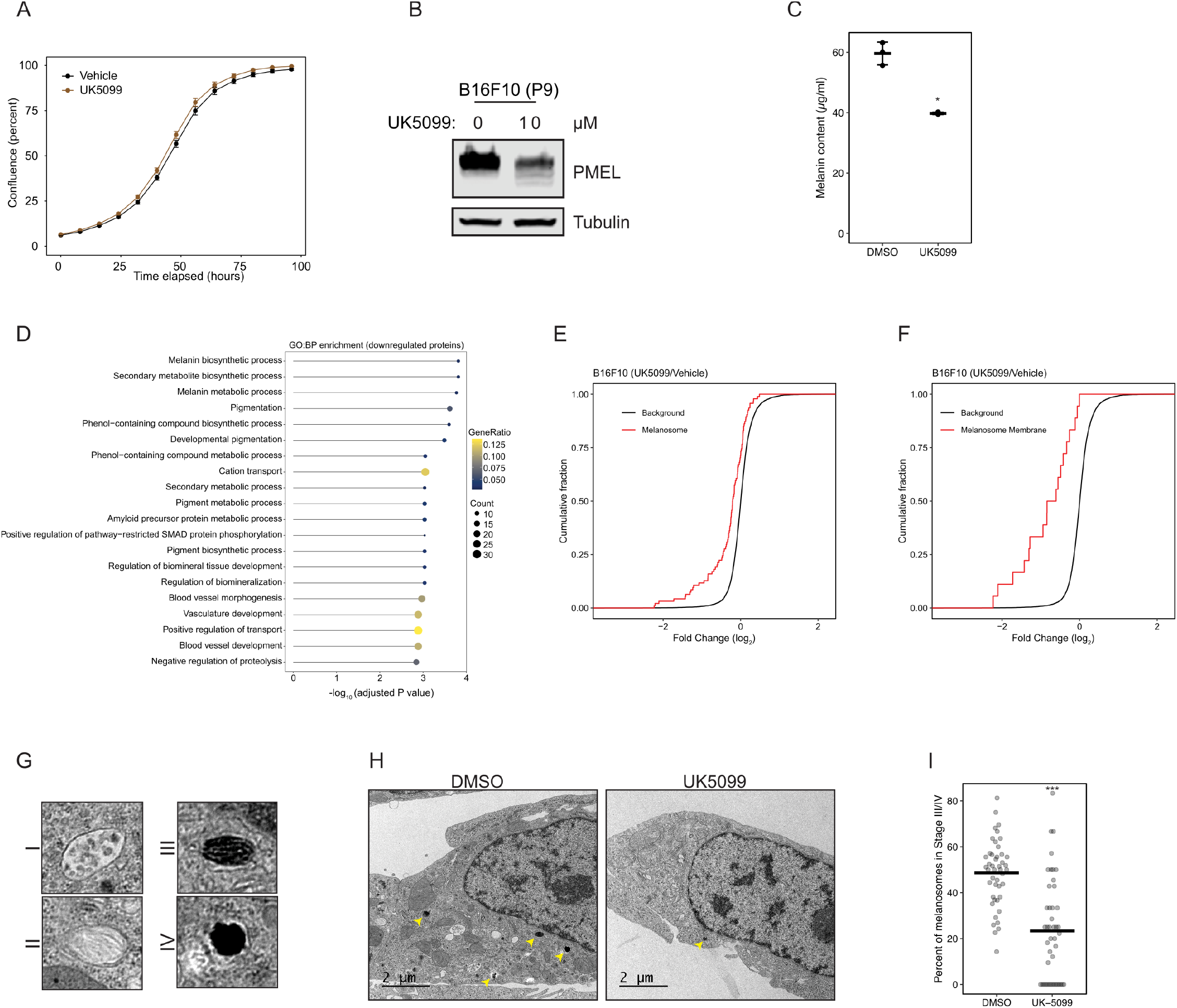
Inhibition of the MPC decreases melanosome protein abundance. **(A)** Line plot showing change in confluence measured by Incucyte of B16F10 cells treated with vehicle or 10 µM UK-5099. **(B)** Immunoblots of B16F10 cells treated with UK5099 for 24 hours at the indicated dose. **(C)** Dot plot showing quantification of melanin content from B16F10 cells treated with vehicle or 10 µM UK-5099 for 48 hours. A two-tailed, two-sample t-test was performed (\**P* < 0.05, \*\**P* < 0.01, \*\*\**P* < 0.001). **(D)** GO term enrichment for biological processes from quantitative proteomics of B16F10 cells treated with UK5099. **(E)** Cumulative distribution plots showing log_2_ fold change in protein abundance comparing UK5099 treated cells to control cells. Genes are annotated as related to the melanosome GO annotation or **(F)** melanosome membrane GO annotation or the background set. A two-tailed Kolmogorov–Smirnov test was performed (*P* = 2.011 × 10^−10^ and *P* = 2.401 × 10^−8^). **(G)** Transmission electron microscopy images of melanosome stages, I-IV as indicated. **(H)** Representative TEM images from B16F10 cells treated with vehicle or 10 µM UK-5099 for 48 hours. Yellow arrows indicate late-stage melanosomes. **(I)** Quantification of the number of late-stage melanosomes from **(H)**. A two-tailed, two-sample t-test was performed (\**P* < 0.05, \*\**P* < 0.01, \*\*\**P* < 0.001).

**Figure S9:**
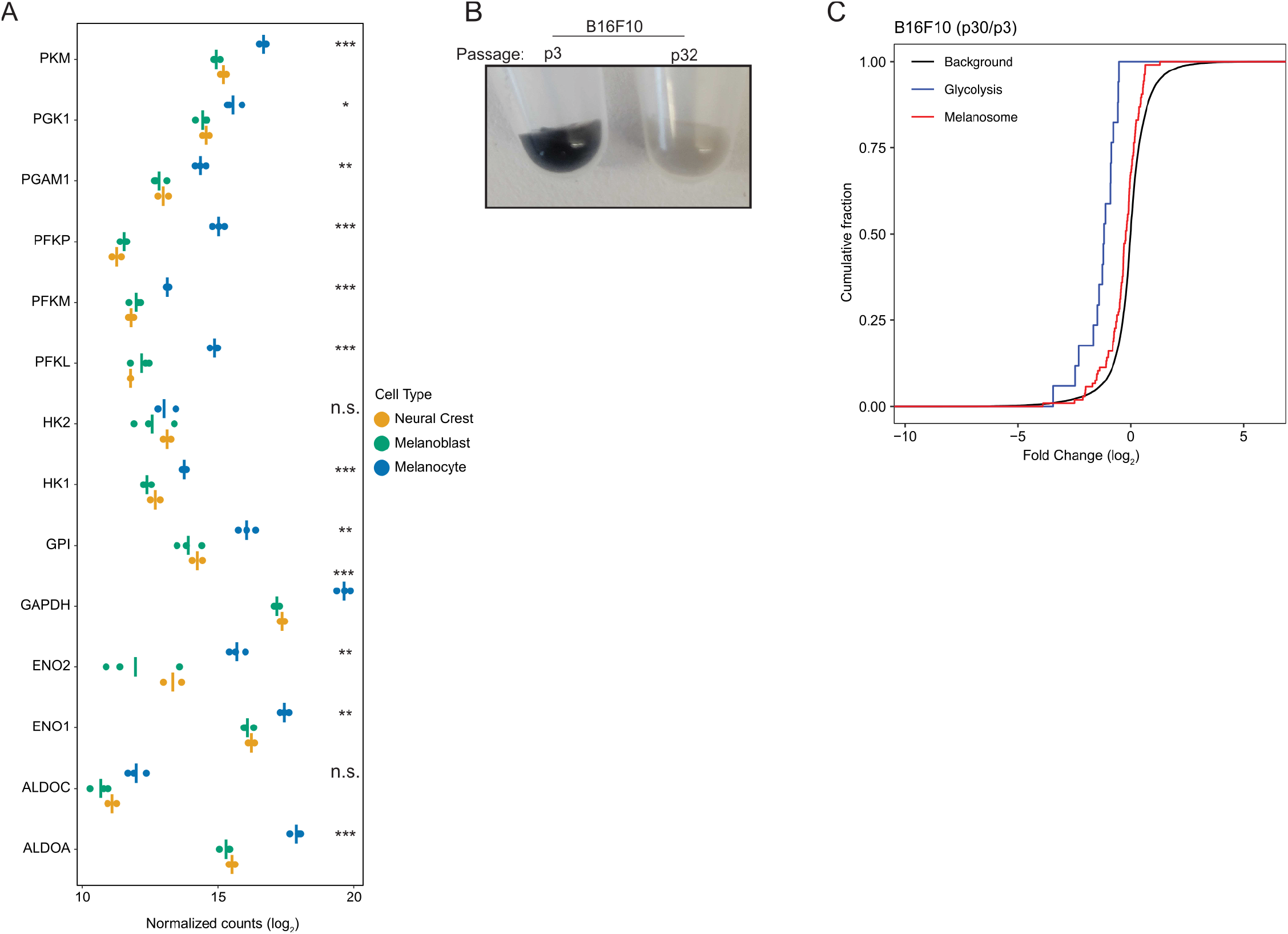
Glycolytic gene expression is correlated with differentiation state and pigmentation. **(A)** Dot plot showing RNA-sequencing normalized counts (GSE172029) of indicated genes from hPSCs differentiated into indicated cell types. A one-way ANOVA and Dunnett test were performed, statistics indicate melanocyte relative to neural crest (n.s. not significant, \**P* < 0.05, \*\**P* < 0.01, \*\*\**P* < 0.001). **(B)** Photograph of B16F10 cells at the indicated passage number. **(C)** Cumulative distribution plots displaying log_2_ fold change comparing late passage to early passage B16F10 cells. Genes are annotated as related to glycolysis, melanosome, or background set. A two-tailed Kolmogorov–Smirnov test was performed between glycolysis and background annotation (*P* = 2.969 × 10^−10^).

**Figure S10:**
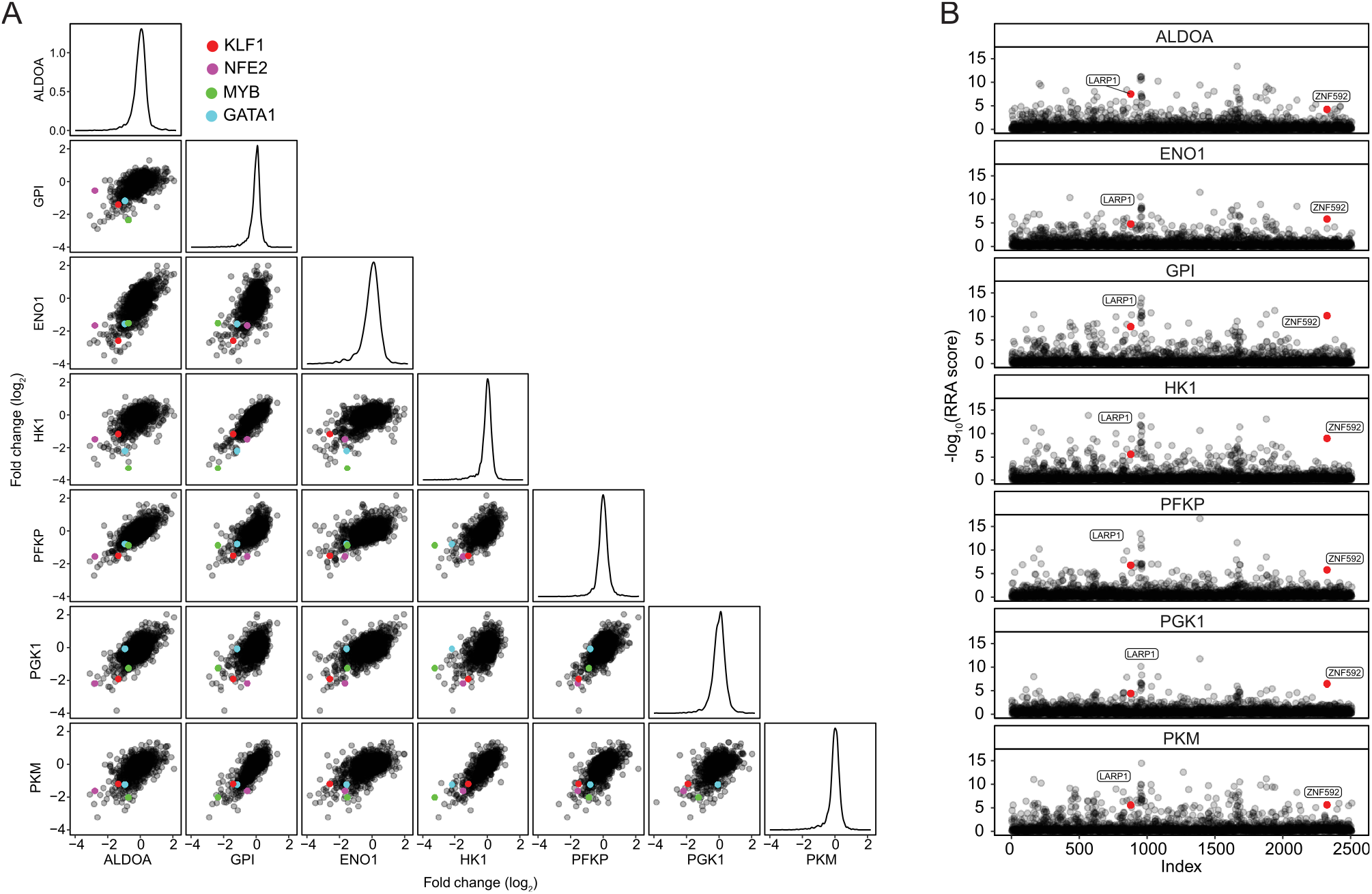
TRoUT-FISH identifies hematopoietic lineage-specifying factors as regulators of glycolysis in K562 cells. **(A)** TRoUT-FISH results across glycolytic genes. Pairwise scatterplots display the gene level log_2_ fold change comparing the high and low populations for each target gene. On the diagonal are density plots displaying the distribution of gene level effects. Red is *KFL1*, magenta is *NFE2*, green is *MYB*, and cyan is *GATA1* in each scatterplot. **(B)** TRoUT-FISH identifies regulators of glycolytic target genes in K562 cells. Scatterplots show genes indexed alphabetically along the x-axis and the RRA scores from MAGeCK comparing the high and low populations of each target gene along the y-axis. *LARP1* and *ZNF592* are labeled and colored in red.

**Figure S11:**
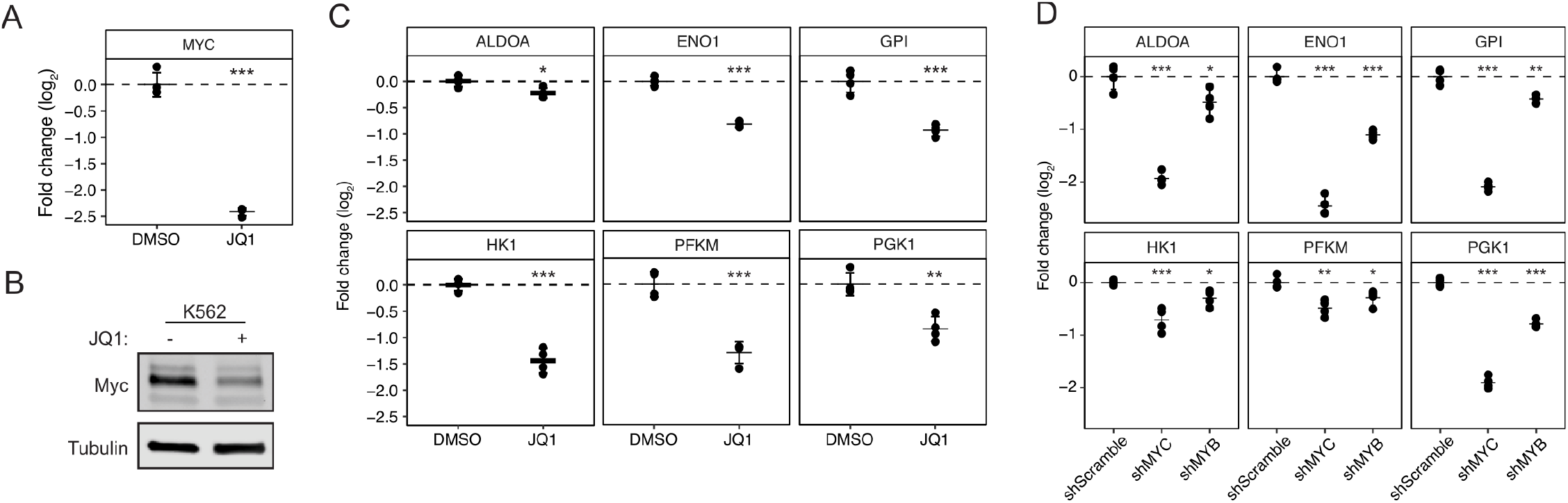
MYC and MYB control metabolic gene expression. **(A)** Dot plot showing changes in MYC expression in K562 cells treated with 1 µM JQ1 for 6h, as determined by qPCR or **(B)** by immunoblot. Experiments performed in quadruplicate and means ± SD is shown. A two-tailed, two-sample t-test was performed (\**P* < 0.05, \*\**P* < 0.01, \*\*\**P* < 0.001). **(C)** Dot plot showing changes in glycolytic gene expression as in **(A)**. Experiments performed in triplicate and means ± SD is shown. A two-tailed, two-sample t-test was performed (\**P* < 0.05, \*\**P* < 0.01, \*\*\**P* < 0.001). **(D)** Dot plot showing changes in glycolytic gene expression K562 cells infected with shRNAs against the indicated genes 5 days post infection, as determined by qPCR. Experiments performed in quadruplicate and means ± SD is shown. A one-way ANOVA and Dunnett test were performed (\**P* < 0.05, \*\**P* < 0.01, \*\*\**P* < 0.001).

**Fig. S12:**
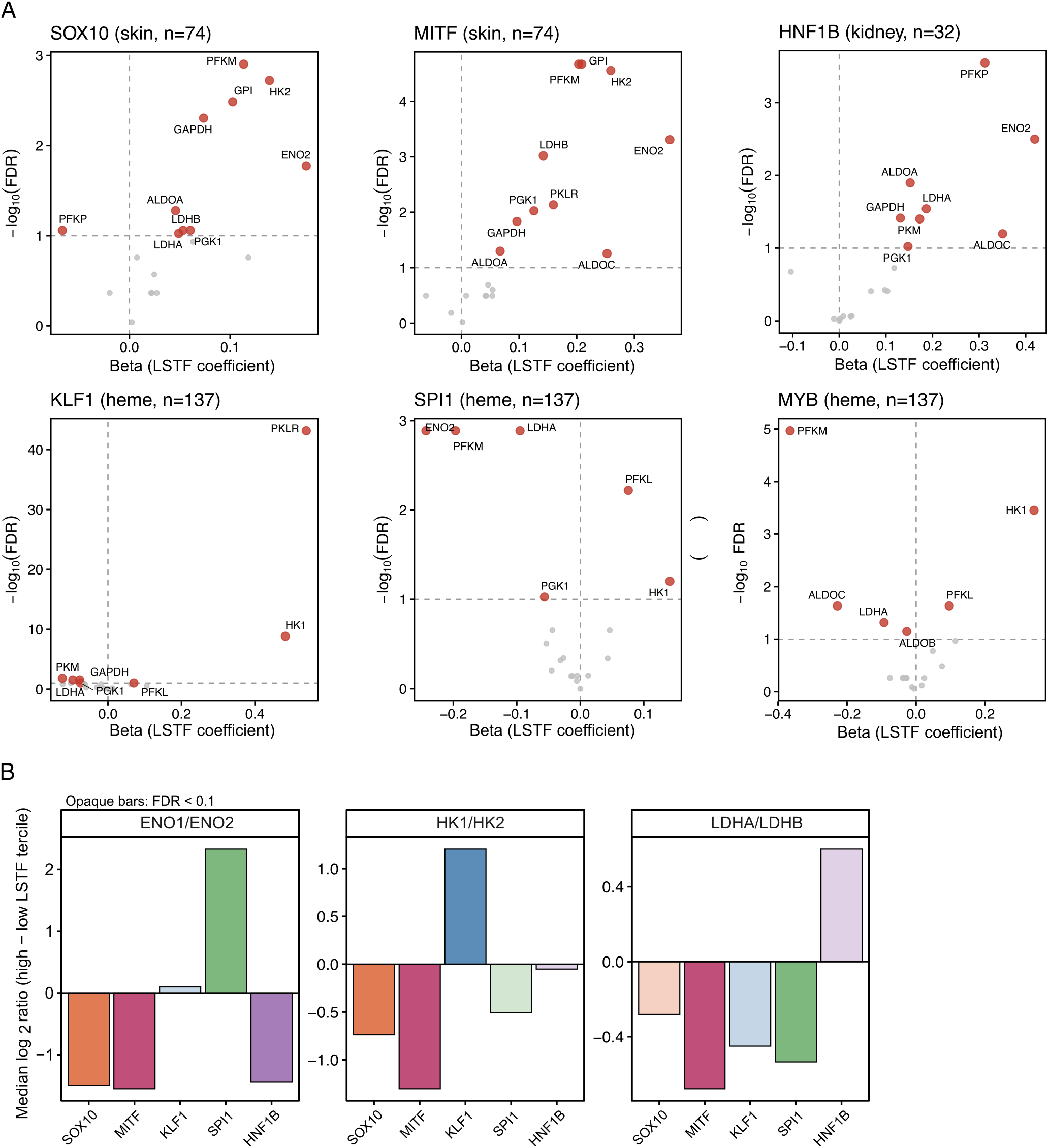
Lineage specifying transcription factors control metabolic gene isoforms. **(A)** Volcano plots showing all 19 glycolytic gene associations for each LSTF from intra-lineage regression (gene ~ LSTF + MYC; see Methods). Each panel represents one LSTF within its matched cancer lineage. Red points with labels indicate genes passing FDR < 0.10; gray points are non-significant. Sample sizes: SOX10 and MITF, skin/melanoma, n = 74; KLF1, SPI1, and MYB, hematopoietic, n = 137; HNF1B, kidney, n = 32. **(B)** Difference in median log_2_ isoenzyme ratio (high LSTF tercile minus low LSTF tercile) for three glycolytic isoenzyme pairs (HK1/HK2, ENO1/ENO2, LDHA/LDHB) within each lineage. A positive value indicates a higher ratio of the first isoform relative to the second in cells with high LSTF expression. Bars are colored by LSTF; opaque bars pass FDR < 0.10 (Wilcoxon test, Benjamini–Hochberg correction); translucent bars are non-significant.

**Fig. S13:**
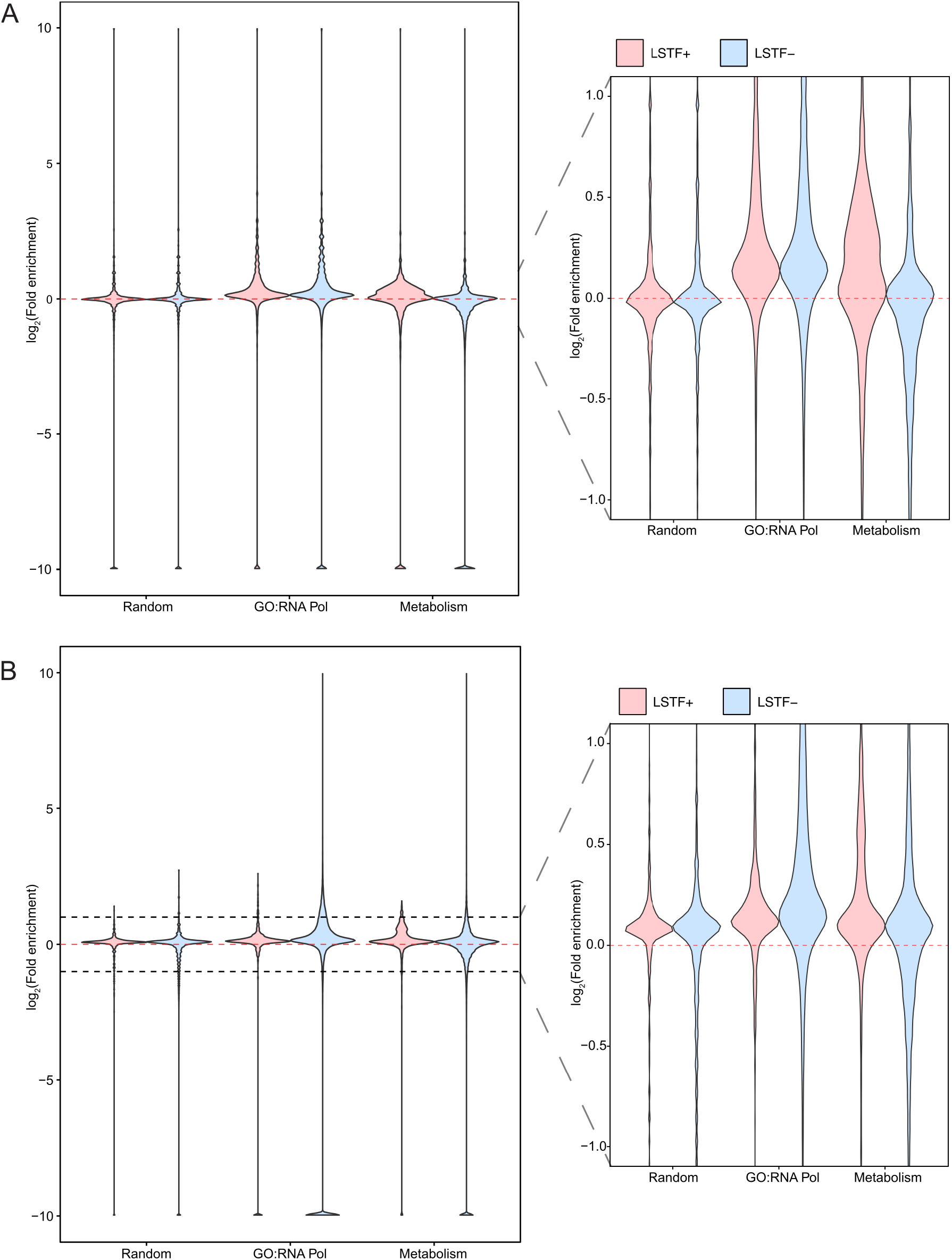
Lineage specifying transcription factors control metabolic gene expression. **(A)** Violin plot showing the log_2_(Fold enrichment) of LSTF binding within 5kb of indicated gene set TSS for human ChIP-Atlas datasets. Insets show the region bounded by log_2_(Fold enrichment) = ± 1. Adjusted odds ratio from logistic regression reported in **(Fig. 4G). (B)** Violin plot showing the log_2_(Fold enrichment) of LSTF binding within 5kb of indicated gene set TSS for mouse ChIP-Atlas datasets. Insets show the region bounded by log_2_(Fold enrichment) = ± 1. Adjusted odds ratio from logistic regression reported in **(Fig. 4G)**.

