## Supplementary material for "Lineage specifying transcription factors determine cell function by direct control of metabolism": DataS3

### BD FACSDiva 8.0

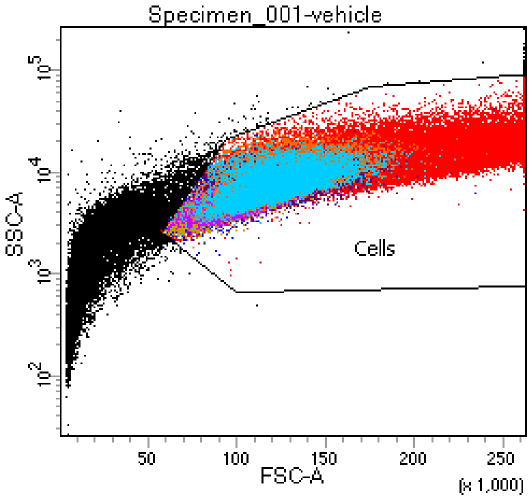


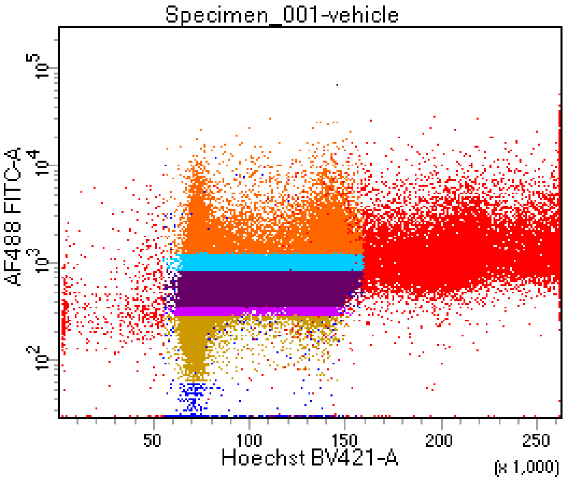


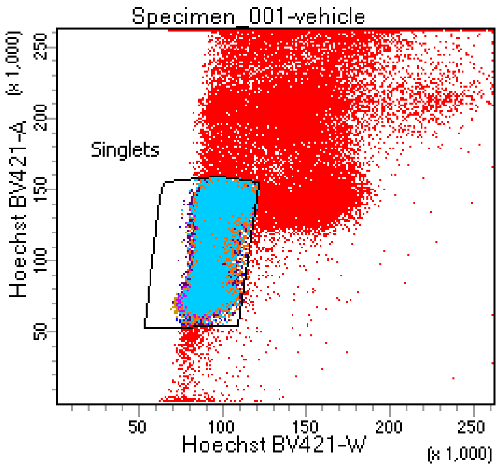


Tube: vehicle

| Population | #Events | %Parent | %Total |
|---|---|---|---|
| All Events | 174,886 | #### | 100.0 |
| Cells | 152,149 | 87.0 | 87.0 |
| Singlets | 108,482 | 71.3 | 62.0 |
| P1 | 46,564 | 42.9 | 26.6 |
| ATTO+ | 106,201 | 97.9 | 60.7 |
| 1 | 13,506 | 12.7 | 7.7 |
| 2 | 12,159 | 11.4 | 7.0 |
| 4 | 9,977 | 9.4 | 5.7 |
| 3 | 10,537 | 9.9 | 6.0 |

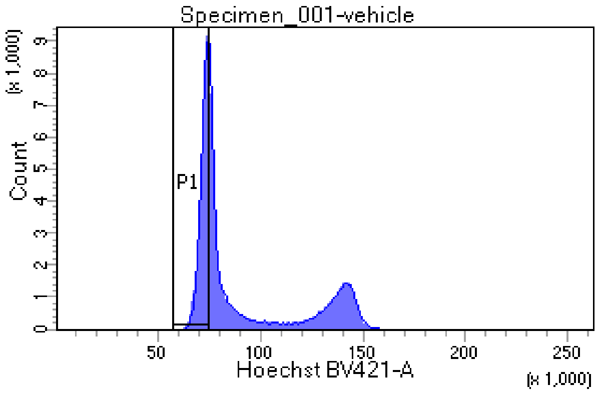


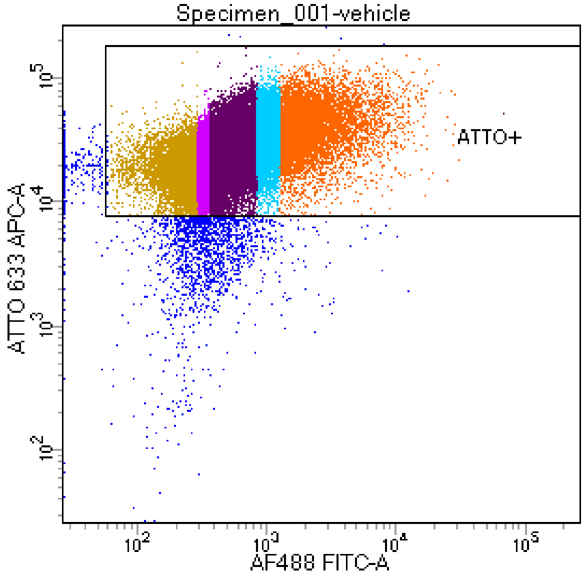

### BD FACSDiva 8.0

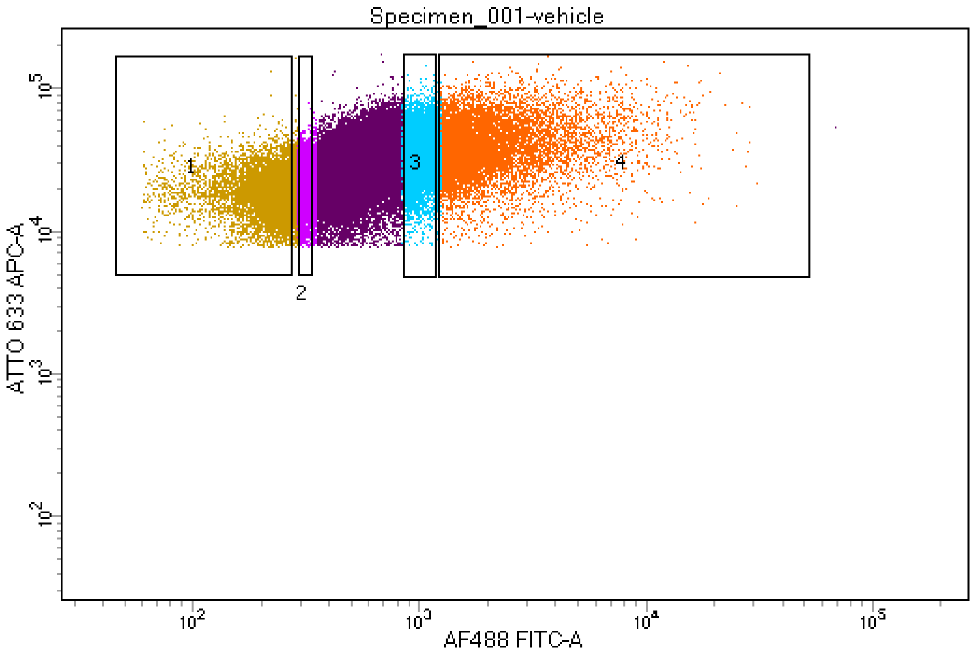


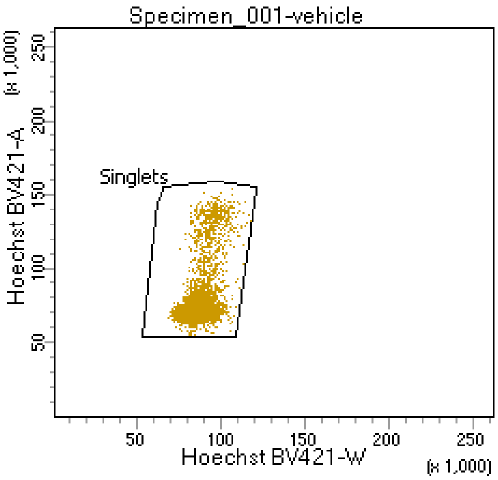


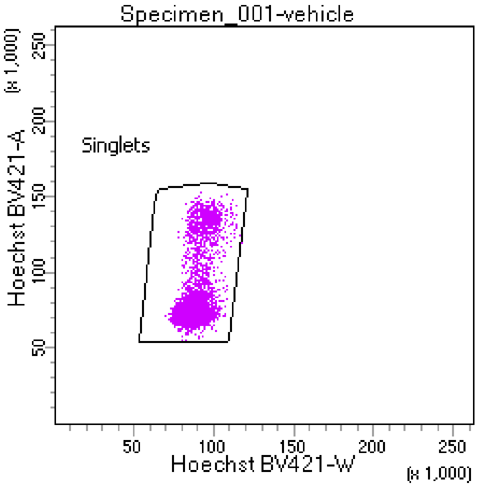


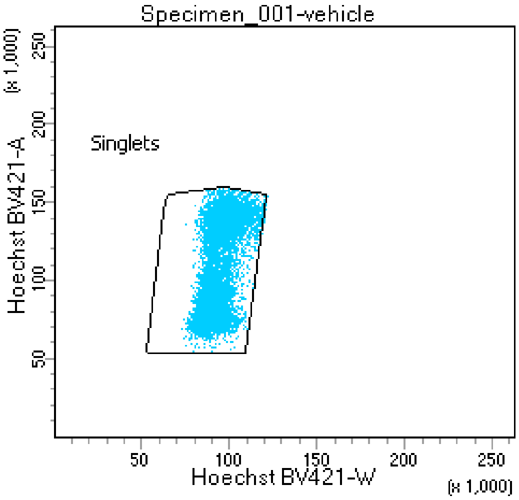


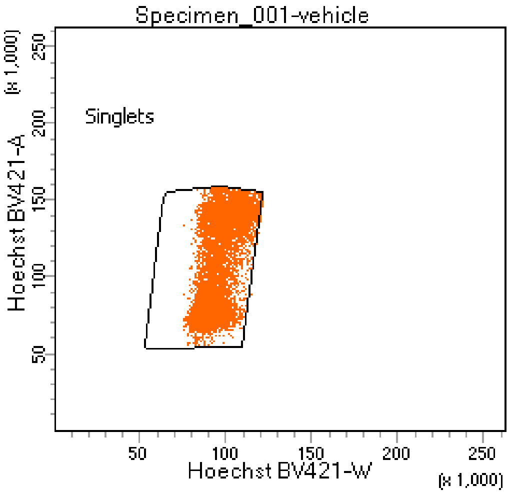

### BD FACSDiva 8.0

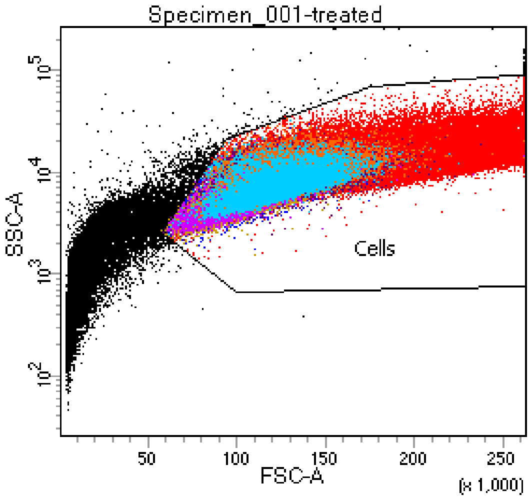


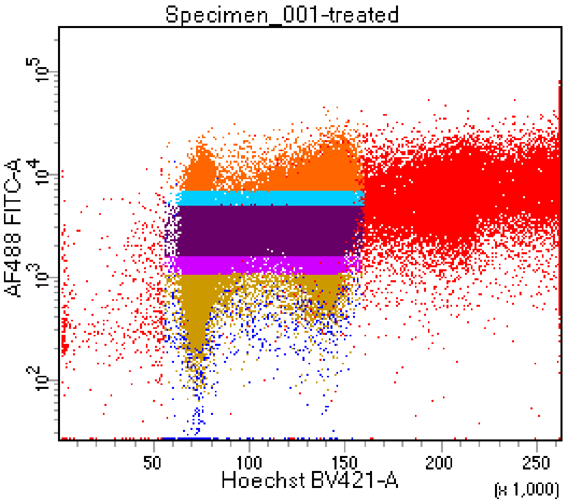


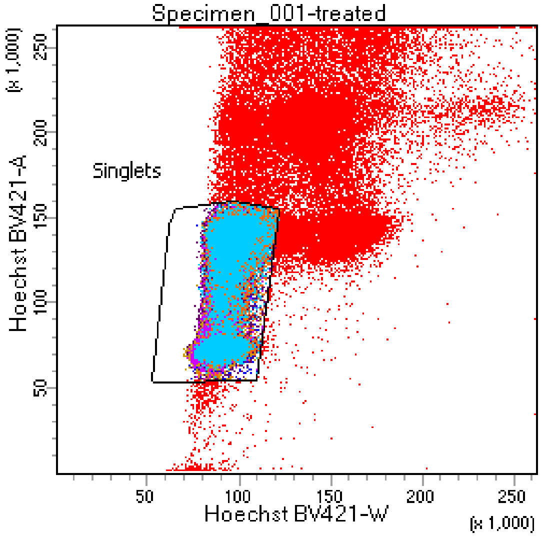


Tube: treated

| Population | #Events | %Parent | %Total |
|---|---|---|---|
| All Events | 184,440 | #### | 100.0 |
| Cells | 164,657 | 89.3 | 89.3 |
| Singlets | 110,826 | 67.3 | 60.1 |
| P1 | 46,218 | 41.7 | 25.1 |
| ATTO+ | 108,471 | 97.9 | 58.8 |
| 1 | 12,345 | 11.4 | 6.7 |
| 2 | 11,762 | 10.8 | 6.4 |
| 4 | 10,969 | 10.1 | 5.9 |
| 3 | 11,154 | 10.3 | 6.0 |

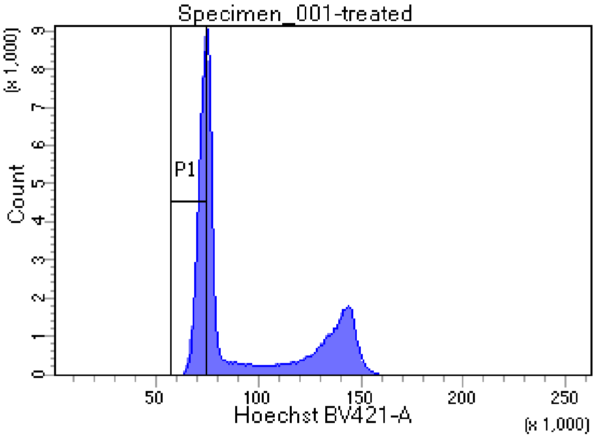


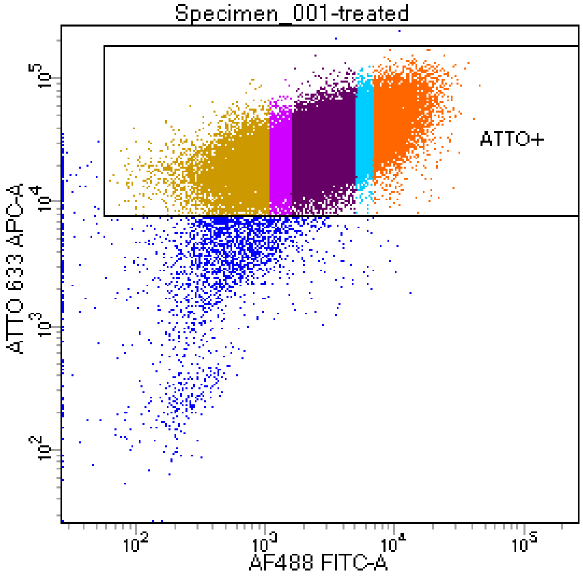

### BD FACSDiva 8.0

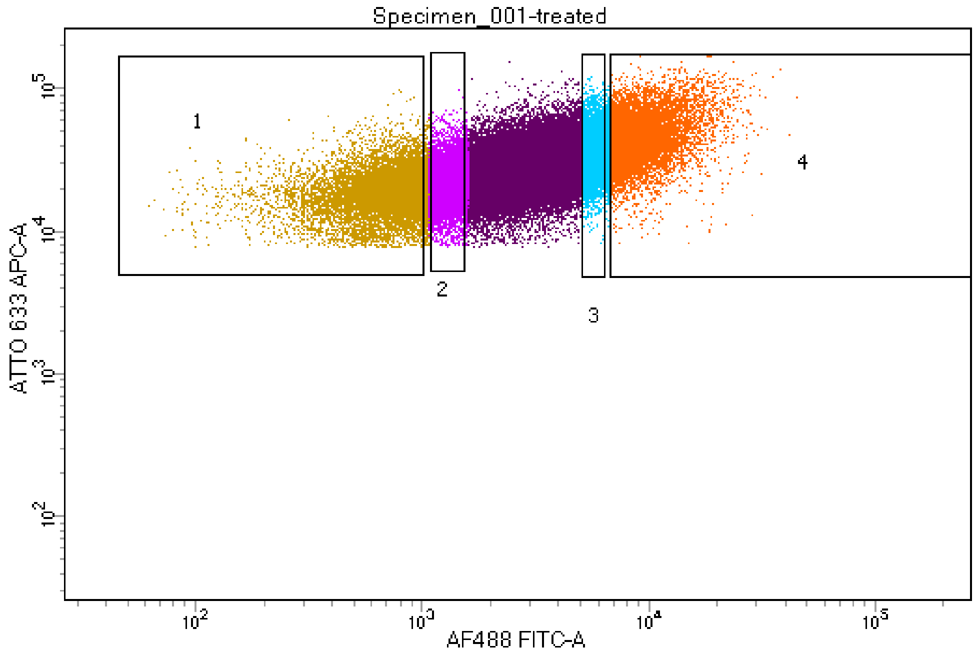


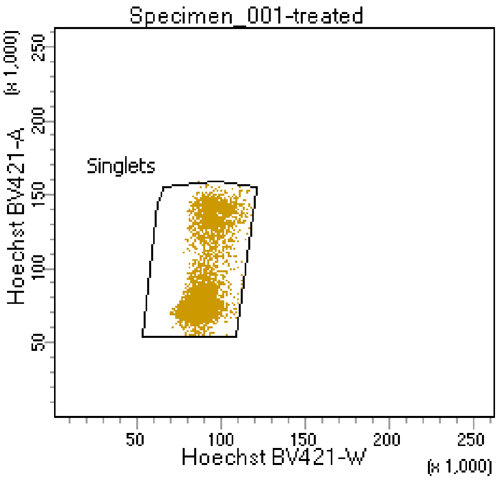


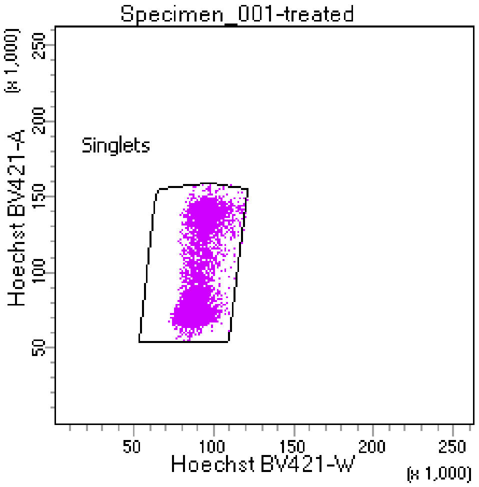


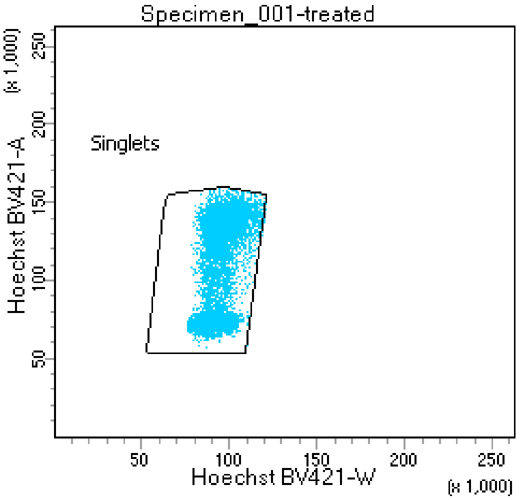


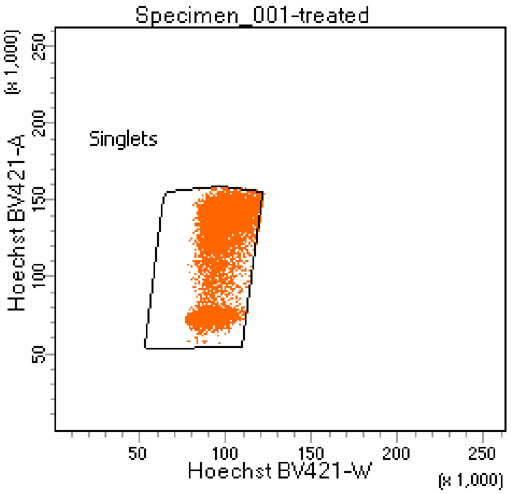
